# A twin UGUA motif directs the balance between gene isoforms through CFIm and the mTORC1 signaling pathway

**DOI:** 10.1101/2022.08.31.506015

**Authors:** R. Samuel Herron, Alexander K. Kunisky, Jessica R. Madden, Vivian I. Anyaeche, Hun-Way Hwang

**Affiliations:** Department of Pathology, University of Pittsburgh, School of Medicine, 3550 Terrace Street, Pittsburgh, PA 15261, USA

**Author notes:** CONTACT INFORMATION: Address: S754 Scaife Hall, 3550 Terrace Street, Pittsburgh, PA, 15261, USA.

## Abstract

Alternative polyadenylation (APA) generates mRNA isoforms and diversifies gene expression. Here we report the identification of a twin UGUA motif, UGUAYUGUA, and its function in APA. Applying cTag-PAPERCLIP to *Tsc1* conditional knockout mice, we discovered that the mTORC1 pathway balances expression of *Trim9* isoforms. We showed that CFIm components, CPSF6 and NUDT21, promote *Trim9/TRIM9-S* expression in mouse and human, and we identified an evolutionarily conserved UGUAYUGUA motif that is critical for this regulation. We found additional CPSF6-regulated polyadenylation sites (PASs) with similar twin UGUA motifs in human, and we experimentally validated the twin UGUA motif functionality in *BMPR1B, MOB4*, and *BRD4-L*. Importantly, we showed that inserting a twin UGUA motif into a heterologous PAS was sufficient to confer regulation by CPSF6 and mTORC1. Our study reveals an evolutionarily conserved mechanism to regulate gene isoform expression and implicates possible gene isoform imbalance in cancer and neurologic disorders with mTORC1 pathway dysregulation.

## INTRODUCTION

Proper expression of gene isoforms is important for normal physiology. For example, in neurons, co-expression of *Cdc42* exon 6 and exon 7 isoforms (both are alternative 3’ terminal exons) is crucial for normal morphogenesis, as loss of either isoform results in abnormal development of axons and dendrites (Yap et al., 2016). APA is an important mechanism to generate RNA isoforms with different 3’ ends (Gruber and Zavolan, 2019; Mitschka and Mayr, 2022; Tian and Manley, 2017). The mRNA 3’-end processing factor CFIm is not essential for the cleavage reaction but has an outsized role in regulating APA as a sequence-dependent activator of mRNA 3’-end processing (Boreikaite et al., 2022; Schmidt et al., 2022; Zhu et al., 2018). The human CFIm consists of two subunits: a small subunit, CFIm25 (encoded by the NUDT21 gene), which directly binds to the UGUA motif, and two alternative large subunits, CFIm68 (encoded by the CPSF6 gene) and CFIm59 (encoded by the CPSF7 gene), which activates 3’-end processing by interacting with CPSF, one of the essential mRNA 3’-end processing factors (Yang et al., 2011a; Zhu et al., 2018). It has been shown that loss of NUDT21 or CPSF6, but not CPSF7, resulted in widespread and overlapping APA alterations in cells (Ghosh et al., 2022; Gruber et al., 2012; Hwang et al., 2016; Li et al., 2015; Martin et al., 2012; Masamha et al., 2014; Zhu et al., 2018). This is consistent with the finding that CFIm59 is a weaker activator of 3’-end processing compared with CFIm68 (Zhu et al., 2018) and highlights the important roles of NUDT21 and CPSF6 in APA regulation. Notably, APA changes from loss of NUDT21 or CPSF6 are predominantly proximal shifts that lead to 3’ UTR shortening or expression of truncated proteins (Gruber et al., 2012; Martin et al., 2012; Masamha et al., 2014), which can be explained by the skewed distribution of the UGUA motif that favors the distal PAS in CFIm target mRNAs (Ghosh et al., 2022; Hwang et al., 2016; Li et al., 2015; Zhu et al., 2018).

The mTORC1 signaling pathway plays a central role in regulating cell metabolism and hyperactive mTORC1 causes a group of neurodevelopmental disorders termed “mTORopathies” with shared clinical manifestations (Crino, 2016; Lipton and Sahin, 2014; Liu and Sabatini, 2020). One of the best-studied mTORopathies is Tuberous Sclerosis Complex (TSC), which is caused by loss-of-function mutations in *TSC1* or *TSC2*, both of which are mTORC1 inhibitors (Salussolia et al., 2019). Interestingly, a recent RNA-seq study identified hundreds of mRNAs with 3’ UTR shortening in *Tsc1*-null mouse embryonic fibroblasts (Chang et al., 2015), which is reminiscent of the aforementioned APA changes from loss of NUDT21 or CPSF6. Subsequently, a direct link between the mTORC1 pathway and CPSF6 was discovered in *Drosophila—*in starvation, repression of the mTORC1 signaling allows two downstream kinases, CDK8 and CLK2, to phosphorylate CPSF6, which is required for its nuclear localization to promote 3’ UTR lengthening of autophagy genes *Atg1* and *Atg8a* (Tang et al., 2018). The regulation of CPSF6 by CDK8 and CLK2 was also present in human MCF7 cells and was required for starvation-induced autophagy (Tang et al., 2018). Taken together, these two studies indicate that CPSF6-mediated APA regulation is a previously underappreciated component of the mTORC1 signaling pathway and suggest that APA dysregulation might contribute to TSC pathogenesis.

Neurons have a highly complex transcriptome that consists of multiple gene isoforms and many long 3’ UTRs not seen in other cell types (Miura et al., 2013; Tushev et al., 2018). Neurologic defects occur when normal APA is disrupted in neurons (Alcott et al., 2020; LaForce et al., 2022). Because primary morbidity for TSC patients comes from the central nervous system involvement (Salussolia et al., 2019), we sought to investigate how hyperactive mTORC1 impacts the neuronal APA landscape in the brain. *Tsc1* conditional knockout mice recapitulate several human disease features and have been widely used for TSC studies (Bateup et al., 2013; Ercan et al., 2017; Kwiatkowski et al., 2002; Meikle et al., 2007). We previously developed the cTag-PAPERCLIP technique, which utilizes a Cre-inducible allele of GFP-tagged poly(A)-binding protein (PABP) to perform high-throughput APA profiling *in vivo* in specific cell types without cell purification in mouse (Hwang et al., 2017). In this study, we applied cTag-PAPERCLIP and adeno-associated virus (AAV) delivery of Cre to knock out *Tsc1* in mouse brain cortical excitatory neurons for *in vivo* APA profiling. We discovered that mTORC1 activities modulate the balance between two *Trim9/TRIM9* isoforms in mouse cortical excitatory neurons *in vivo* and in differentiated human neural stem cells (NSCs) in culture. *Trim9/TRIM9* encodes a neuronally enriched E3 ubiquitin ligase that regulates neuron morphogenesis (Winkle et al., 2014; 2016). We found that expression of the short *Trim9/TRIM9* isoform was dependent on CPSF6 and NUDT21, and we identified an evolutionarily conserved twin UGUA motif (UGUAYUGUA) that is essential for its PAS usage. We further demonstrated the existence of similar functional twin-UGUA motifs in additional human CPSF6-dependent PASs. Importantly, we showed that it is possible to engineer a PAS to be regulated by CPSF6 and mTORC1 by insertion of a twin UGUA motif. Overall, our study identifies an evolutionarily conserved mechanism to regulate gene isoform expression by the mTORC1 pathway and expands current knowledge of APA regulation by CFIm beyond the well-established UGUA motif.

## RESULTS

### Systemic AAV delivery of Cre recombinase is an efficient way to model human disease in the cTag-PABP mouse for APA profiling

In our original cTag-PAPERCLIP study (Hwang et al., 2017), the APA profiling was performed on double-heterozygous mice with 2 transgenes: a cell type-specific Cre recombinase (from a Cre driver mouse) and the Cre-inducible allele of GFP-tagged PABP (PABP-GFP; from the cTag-PABP mouse). In order to conditionally knock out *Tsc1* for our current study, a third transgene—the floxed *Tsc1* allele from the *Tsc1*-floxed mice—has to be present in homozygosity, and we would like to avoid the inefficient and time-consuming breeding process to generate mice with 3 transgenes. Recent development of the PHP.eB capsid makes it possible to broadly and efficiently transduce brain neurons from systemic AAV injection (Chan et al., 2017). Therefore, we sought to eliminate genetic breeding with Cre driver mice and deliver Cre through systemic AAV-PHP.eB injection instead (Fig. 1A and Fig. S1A).

**Figure 1.**
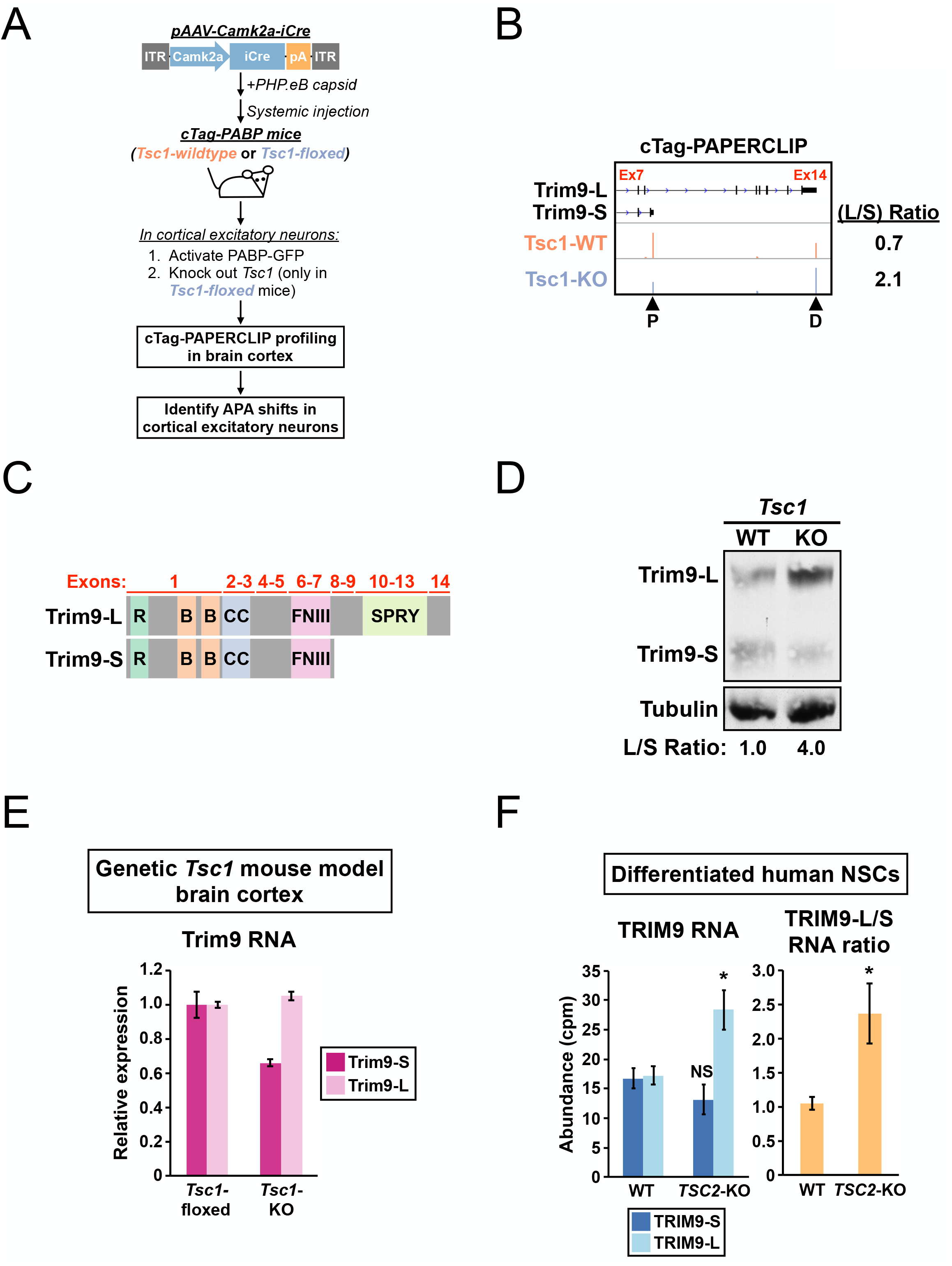
A shift toward Trim9/TRIM9-L expression in *Tsc1-KO* mouse cortical excitatory neurons and differentiated human *TSC2-KO* NSCs. (**A**) The experimental strategy to identify *in vivo* APA shifts in *Tsc1-KO* cortical excitatory neurons in mouse. (**B**) GENCODE annotations and cTag-PAPERCLIP results (merged from two biological replicates) for *Trim9*. Arrowheads: poly(A) sites identified by cTag-PAPERCLIP. P: Proximal. D: Distal. (**C**) Illustrations comparing the exons and known protein domains (based on UniProt annotations) contained in mouse Trim9-L and Trim9-S. R, RING-type Zinc finger. B, B box-type Zinc finger. CC, Coiled coil. FNIII, Fibronectin type-III. (**D**) Western blots showing a shift toward Trim9-L protein expression in a pAAV-Camk2a-iCre injected *Tsc1^fl/fl^*; cTag-PABP mouse (KO) in the brain cortex when compared to an uninjected cTag-PABP mouse (WT). Tubulin: loading control. (**E**) Quantitation of *Trim9* mRNA isoforms by RT-qPCR in the brain cortex of *Tsc1^fl/fl^* (“*Tsc1*-floxed”) and *Camk2a-Cre; Tsc1^fl/fl^* (“*Tsc1*-KO”) mice (the same pair of mice shown in Fig. S1E). (**F**) Quantitation of individual *TRIM9* mRNA isoforms (left) or TRIM9-L/TRIM9-S mRNA ratio (right) by RNA-seq (GSE78961) in *TSC2*-wildtype (WT) and *TSC2*-KO human NSCs after 6 weeks of neural differentiation. cpm: counts per million. Error bars indicate SEM. Statistical significance is determined by two-tailed t-test (1F, left panel) and one-tailed t-test (1F, right panel). NS: not significant, *: p<0.05.

To test the feasibility of the planned AAV delivery strategy, we chose the excitatory neuron-specific mouse *Camk2a* promoter (Madisen et al., 2010) to generate a prototype AAV-Cre, pAAV-Camk2a-iCre. We selected the *Camk2a* promoter for two reasons: First, we have extensive experience using *Camk2a-Cre* mice for cTag-PAPERCLIP profiling (Hwang et al., 2017). Second, knocking out *Tsc1* in excitatory neurons of *Camk2a-Cre; Tsc1^fl/fl^* mice results in TSC-relevant phenotypes (Bateup et al., 2013). Next, we injected wildtype (WT) cTag-PABP mice with pAAV-Camk2a-iCre at two different doses and performed cTag-PAPERCLIP using brain cortex tissues to assess the efficiency of Cre delivery 3 weeks after injection. A genetically bred *Camk2a-Cre;* cTag-PABP mouse was included in the cTag-PAPERCLIP experiment for comparison. On autoradiography, all 3 mice had similar intensities of radioactive signals from the immunoprecipitated PABP-GFP:RNA complexes (Fig. S1B), suggesting that Cre expression through systemic AAV delivery is as efficient as genetic breeding to Cre driver mice for cTag-PAPERCLIP profiling.

Next, we proceeded to use pAAV-Camk2a-iCre to delete *Tsc1* in brain cortical excitatory neurons in cTag-PABP mice. We injected adult *Tsc1*-WT (*Tsc1^+/+^*) and *Tsc1*-floxed (*Tsc1*^fl/fl^) cTag-PABP mice with pAAV-Camk2a-iCre. We sacrificed the injected mice 2~3 weeks after injection and performed cTag-PAPERCLIP profiling to identify APA shifts from activation of the mTORC1 signaling in brain cortical excitatory neurons *in vivo* (Fig. 1A). We expected pAAV-Camk2a-iCre injection to: 1) turn on PABP-GFP expression in excitatory neurons for cTag-PAPERCLIP profiling in both *Tsc1*-WT and *Tsc1*-floxed cTag-PABP mice; 2) knock *Tsc1* out and activate the mTORC1 signaling in excitatory neurons in *Tsc1*-floxed cTag-PABP mice. To verify the expected effects from pAAV-Camk2a-iCre injection, we first examined *Tsc1* expression in the cTag-PAPERCLIP profiles. As expected, *Tsc1* mRNA expression was strongly reduced (>80%, 44.9 to 8.6 cpm, p<0.01) in the injected *Tsc1*-floxed cTag-PABP mice (designated as *Tsc1-KO* hereafter) while both *Tsc2* and *Mtor* mRNAs were expressed at similar levels in the injected mice of both genotypes (Fig. S1C). Next, we checked whether the decrease in *Tsc1* expression was sufficient to activate mTORC1 signaling in the *Tsc1*-KO cTag-PABP mice. We performed western blots for both total and phosphorylated S6 (PS6) ribosomal protein, a well-established indicator of mTORC1 activities (Sengupta et al., 2010). As expected, PS6 in brain cortex was strongly increased in an injected *Tsc1*-floxed cTag-PABP mouse when compared to an uninjected mouse of the same genotype (Fig. S1D). Importantly, the increase in PS6 between the injected-uninjected pair of *Tsc1*-floxed cTag-PABP mice is identical to the increase in PS6 between a *Camk2a-Cre; Tsc1^fl/fl^* mouse and a *Tsc1^fl/fl^* mouse (Fig. S1E), indicating that Cre delivery efficiency from systemic AAV injection is equivalent to that of genetic breeding. Therefore, we concluded that systemic AAV injection is an effective way to deliver cell type-specific Cre recombinases for cTag-PAPERCLIP profiling.

### A shift toward Trim9/TRIM9-L expression in *Tsc1-KO* mouse brain and differentiated human *TSC2*-KO NSCs

From the cTag-PAPERCLIP experiments (two biological replicates for each genotype), we identified 135 genes with 2 PASs that significantly changed their APA preference (FDR<0.05, >2-fold change) in *Tsc1-KO* cortical excitatory neurons—30 shifted proximally while 105 shifted distally (Table S1). To select candidate genes for further investigation, we searched for APA shifts that satisfied the following criteria (Kunisky et al., 2021): 1) switched the major PAS, defined as the predominantly used (>50% of total read counts) PAS between two PASs, and 2) resulted in expression of different protein isoforms. Only one gene from the list, *Trim9*, fulfilled both criteria. *Trim9* encodes a neuronally enriched E3 ubiquitin ligase that regulates neuron morphogenesis (Winkle et al., 2014; 2016). The two *Trim9* isoforms identified by cTag-PAPERCLIP in mouse cortical excitatory neurons are designated as Trim9-L and Trim9-S hereafter. Trim9-L is the full-length isoform while Trim9-S uses an upstream PAS to skip exons 8 to 14 and lacks the SPRY domain at the protein level (Fig. 1B and 1C).

We next performed western blotting with brain cortices from *Tsc1-WT* and *Tsc1*-KO cTag-PABP mice to examine the impact of *Trim9* APA shift at the protein level. In parallel with the cTag-PAPERCLIP results, a shift toward Trim9-L protein expression was also observed in the *Tsc1*-KO cTag-PABP mouse (Fig. 1D). To exclude the possibility that the observed *Trim9* APA shift is specific to pAAV-Camk2a-iCre injected cTag-PABP mice, we measured expression of Trim9-L and Trim9-S mRNA isoforms by RT-qPCR using brain cortex tissues from the *Camk2a-Cre; Tsc1^fl/fl^* mouse and *Tsc1^fl/fl^* mouse pair shown in Fig. S1E. Although this assay is not cell-type specific like cTag-PAPERCLIP, a similar shift toward Trim9-L mRNA expression was also observed (Fig. 1E). Lastly, because the *Trim9* exon arrangement is conserved in human, we sought to determine whether activation of mTORC1 signaling also favors TRIM9-L expression in human cells of neuronal lineage. We measured the abundance of TRIM9-L and TRIM9-S isoforms in a published RNA-seq dataset generated from WT and *TSC2*-KO human NSCs after 6 weeks of neural differentiation (Grabole et al., 2016). We found that TRIM9-L mRNA expression was increased by 1.7-fold in *TSC2*-KO NSCs compared to WT NSCs (Fig. 1F, left panel). Moreover, the TRM9-L/TRIM9-S ratio was consistently higher in *TSC2*-KO NSCs compared to WT NSCs in all replicates (Fig. 1F, right panel). Taken together, these results show that, in both mouse and human neurons, hyperactive mTORC1 causes an APA shift in *Trim9/TRIM9* that favors the full-length Trim9/TRIM9-L expression.

### The mTORC1 signaling pathway regulates *Trim9/TRIM9* isoform expression in mouse and human cells

We next sought to establish cell culture models to study the mechanistic link between the mTORC1 signaling pathway and *Trim9/TRIM9* APA in human and mouse. We first examined whether manipulation of mTORC1 activities in mouse Neuro-2a (N2a) cells would recapitulate the *Trim9* APA shift observed in mouse cortical neurons *in vivo*. Since mTORC1 is active under normal growth conditions in culture, we started by treating N2a cells with Torin 1, a potent mTORC1 inhibitor (Thoreen et al., 2012), and examined expression of Trim9-L and Trim9-S by RT-qPCR and western blotting. As expected, Torin 1 treatment successfully shut down mTORC1 and eliminated PS6 in N2a cells (Fig. 2A). As predicted from our *in vivo* findings (low mTORC1 activities would favor Trim9-S expression), in Torin 1-treated N2a cells, *Trim9* expression was consistently shifted toward Trim9-S with similar magnitudes at both mRNA (Fig. 2B) and protein (Fig. S2A and Fig. 2C-D) levels. Next, we went the opposite direction and evaluated whether we could further elevate mTORC1 activities in N2a cells by knocking down *Tsc1* or *Tsc2* using siRNAs (Fig. S2B). As evidenced by a rise in PS6 (Fig. 2E), we found that either siTsc1 or siTsc2 transfection could increase mTORC1 activities in N2a cells, which then favored Trim9-L mRNA expression as expected (Fig. 2F).

**Figure 2.**
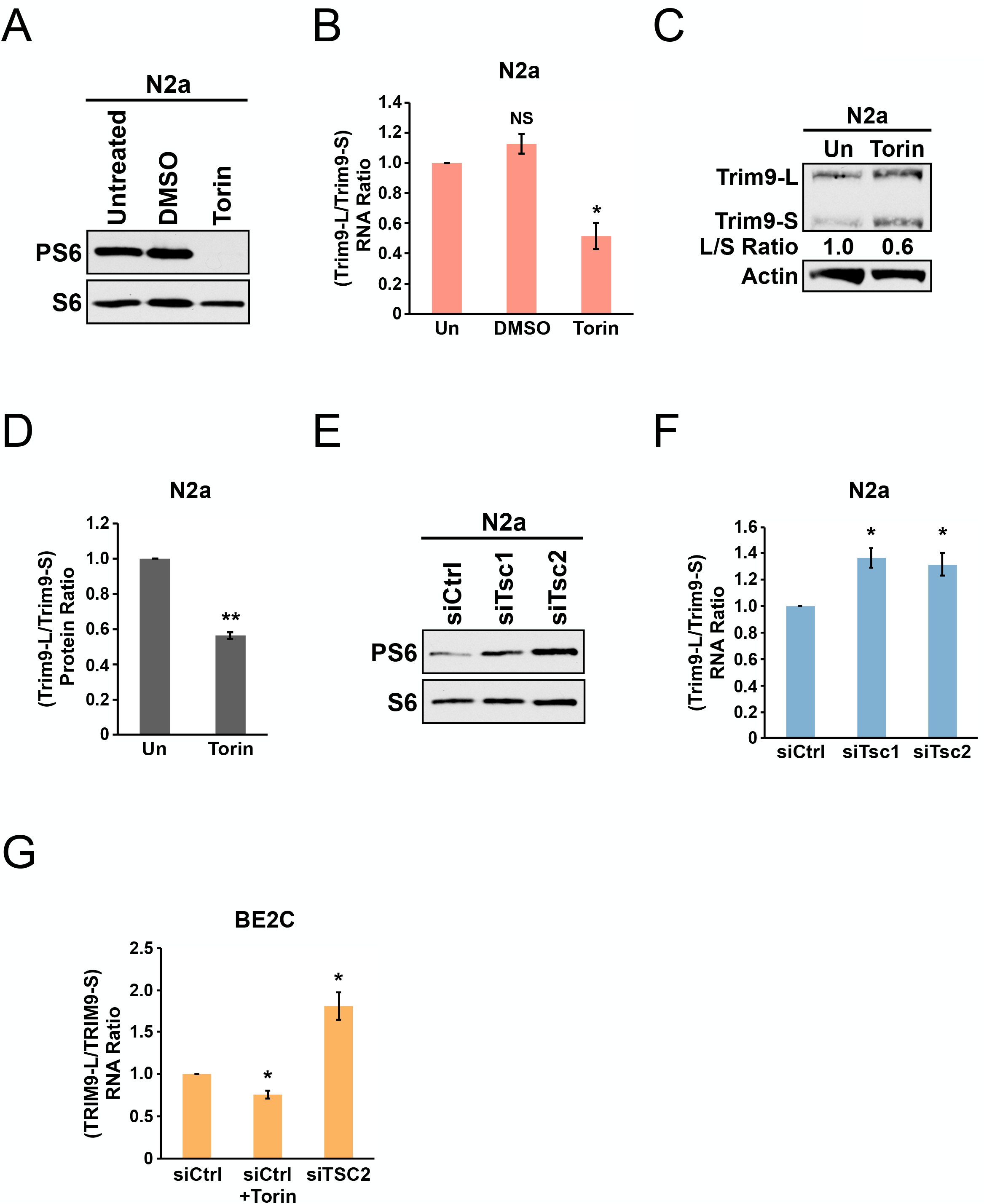
The mTORC1 signaling pathway regulates the balance between Trim9/TRIM9-L and Trim9/TRIM9-S in mouse and human cells. (**A**) Western blots showing expression of total and phosphorylated S6 (PS6) ribosomal protein in N2a cells with different treatments for 48 h. (**B**) Bar graphs showing the ratio of *Trim9* mRNA isoforms measured by RT-qPCR in N2a cells receiving different treatments for 48 h from 4 independent experiments (n=4). (**C**) Western blots showing expression of *Trim9* protein isoforms in untreated (Un) or Torin 1-treated (for 48 h) N2a cells. Actin: loading control. (**D**) Bar graphs showing the ratio of *Trim9* protein isoforms measured by western blotting in untreated (Un) or Torin 1-treated (for 48 h) N2a cells from 3 independent experiments (n=3). (**E**) Western blots showing expression of total and phosphorylated S6 ribosomal protein in N2a cells transfected with different siRNAs for 72 h. siCtrl: control siRNA. (**F**) Bar graphs showing the ratio of *Trim9* mRNA isoforms measured by RT-qPCR in N2a cells transfected with different siRNAs for 72 h from 4 independent experiments (n=4). (**G**) Bar graphs showing the ratio of *TRIM9* mRNA isoforms measured by RT-qPCR in BE2C cells receiving different treatments (siCtrl+Torin: 48 h; siCtrl and siTSC2: 72 h) from 4 independent experiments (n=4). Torin 1 was used at 250nM in all experiments. Due to different exposure conditions, PS6 and S6 levels cannot be directly compared between Fig. 2A and Fig. 2E. Error bars indicate SEM. Statistical significance is determined by two-tailed t-test. NS: not significant, *: p<0.05; **, p<0.01.

Next, we evaluated human BE2C neuroblastoma cells, which can be differentiated into neuron-like cells and are a good host for siRNA transfection (Ogorodnikov et al., 2018). We measured TRIM9-L and TRIM9-S mRNA expression under 3 different conditions: control siRNA transfection (baseline), control siRNA transfection plus Torin 1 treatment (low mTORC1 activities), and *TSC2* siRNA transfection (high mTORC1 activities). As seen in mouse cortical neurons *in vivo* and in N2a cells, low mTORC1 activities indeed promoted TRIM9-S expression while high mTORC1 activities favored TRIM9-L expression in BE2C cells (Fig. 2G and Fig. S2C). Taken together, our cell culture studies recapitulated the *Trim9* APA shift from *Tsc1*-KO mice and showed that the mTORC1 signaling pathway controls the balance between the two *Trim9/TRIM9* isoforms in both mouse and human.

### CPSF6 and NUDT21 promote Trim9/TRIM9-S expression in mouse and human cells

Next, we wished to identify APA factor(s) that regulate *Trim9* APA. We first investigated a possible role of CFIm because mTORC1 signaling has been shown to modulate APA of autophagy genes through CPSF6 in *Drosophila* (Fig. S3)(Tang et al., 2018). We performed CRISPR gene editing to generate N2a cells with *Cpsf6* loss-of-function (Fig. 3A). Interestingly, loss of *Cpsf6* had different effects on the two *Trim9* isoforms: it strongly decreased the abundance of Trim9-S mRNA but did not change Trim9-L mRNA expression (Fig. 3B). Next, we induce *Nudt21* loss-of-function in N2a cells by siRNA transfection (Fig. 3C), which phenocopied loss of *Cpsf6:* Trim9-S mRNA expression was decreased but Trim9-L mRNA expression did not statistically significantly change (Fig. 3D). Lastly, we knocked down *CPSF6* and *NUDT21* separately by siRNAs in BE2C cells (Fig. 3E). Both treatments similarly lowered TRIM9-S expression (Fig. 3F). Taken together, these results suggest that both CPSF6 and NUDT21 promote Trim9/TRIM9-S expression in mouse and human cells.

**Figure 3.**
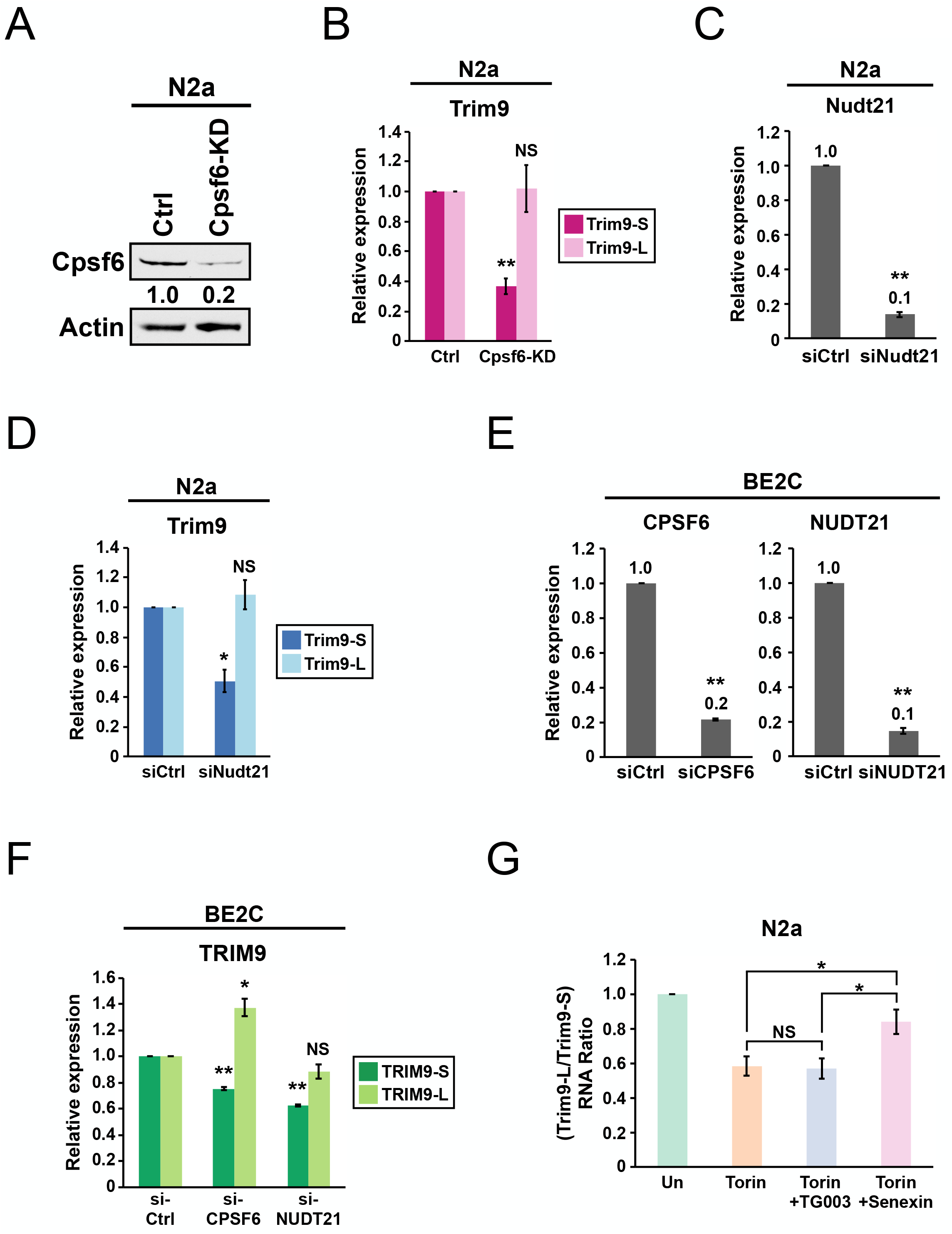
CPSF6 and NUDT21 promote Trim9/TRIM9-S expression in mouse and human cells. (**A**) Western blots showing *Cpsf6* protein expression in control (Ctrl) and *Cpsf6* knockdown (Cpsf6-KD) N2a cells. Actin: loading control. (**B**) Bar graphs showing expression of *Trim9* mRNA isoforms measured by RT-qPCR in Ctrl and Cpsf6-KD N2a cells from 3 independent experiments (n=3). (**C**) Bar graphs showing knockdown efficiency of *Nudt21* siRNAs (siNudt21) measured 72 h after transfection by RT-qPCR from 3 independent experiments (n=3). siCtrl: control siRNA. (**D**) Bar graphs showing expression of *Trim9* mRNA isoforms measured 72 h after transfection by RT-qPCR in N2a cells from the same experiments in (C) (n=3). (**E**) Bar graphs showing knockdown efficiency of *CPSF6* (siCPSF6) and *NUDT21* (siNUDT21) siRNAs measured 72 h after transfection by RT-qPCR from 3 independent experiments (n=3). (**F**) Bar graphs showing expression of *TRIM9* mRNA isoforms measured 72 h after transfection by RT-qPCR in BE2C cells from the same experiments in (E) (n=3). (**G**) Bar graphs showing the ratio of *Trim9* mRNA isoforms measured by RT-qPCR in N2a cells receiving different treatments for 48 h from 4 independent experiments (n=4). Torin 1: 250nM. TG003 and Senexin A: 50μM. Error bars indicate SEM. Statistical significance is determined by two-tailed t-test. NS: not significant, *: p<0.05, **: p<0.01.

In *Drospophila*, CDK8 and CLK2 phosphorylate CPSF6 to promote APA of autophagy genes (Fig. S3)(Tang et al., 2018). Therefore, we asked whether CDK8 and CLK2 also participate in the regulation of *Trim9* APA by mTORC1. In the previous section, we showed that Torin 1 inhibition of mTORC1 favored Trim9-S expression in N2a cells (Fig. 2B). Therefore, we examined whether adding a CDK8 inhibitor or a CLK kinase inhibitor could reverse Torin 1-mediated APA shift toward Trim9-S in N2a cells. We found that TG003 (a CLK kinase inhibitor) did not affect the balance between *Trim9* mRNA isoforms (Fig. 3G). In contrast, Senexin A (a CDK8 inhibitor) partially reversed the shift toward Trim9-S (Fig. 3G). There results suggest that CDK8, but not CLK2, participates in the regulation of *Trim9* APA shift in N2a cells.

### CPSF6 regulates TRIM9-S expression through an evolutionarily conserved twin UGUA motif

Because CFIm binds to the UGUA motif to promote mRNA 3’-end processing (Yang et al., 2011a; Zhu et al., 2018), we inspected the nucleotide sequence of mouse Trim9-S PAS and human TRIM9-S PAS for UGUA motifs. Both mouse Trim9-S PAS and human TRIM9-S PAS contain a 5’ twin UGUA motif (UGUAYUGUA; Y=C in human and T in mouse) followed by two downstream UGUA motifs, for a total of 4 UGUA motifs (Fig. 4A and Fig. S4A). Interestingly, 3 of the 4 UGUA motifs, including the twin UGUA motif, were evolutionarily conserved between mouse and human. To test the biological effects of the identified UGUA motifs on TRIM9-S PAS usage, we developed a tandem PAS reporter assay (Fig. 4B). In this assay, we inserted the PAS of interest into the reporter upstream of the bovine growth hormone (bGH) PAS (which does not contain any UGUA motif), and we measured the abundance of mRNA isoforms generated from both PASs separately by RT-qPCR using PAS-specific primers to infer the relative usage of the PAS of interest.

**Figure 4.**
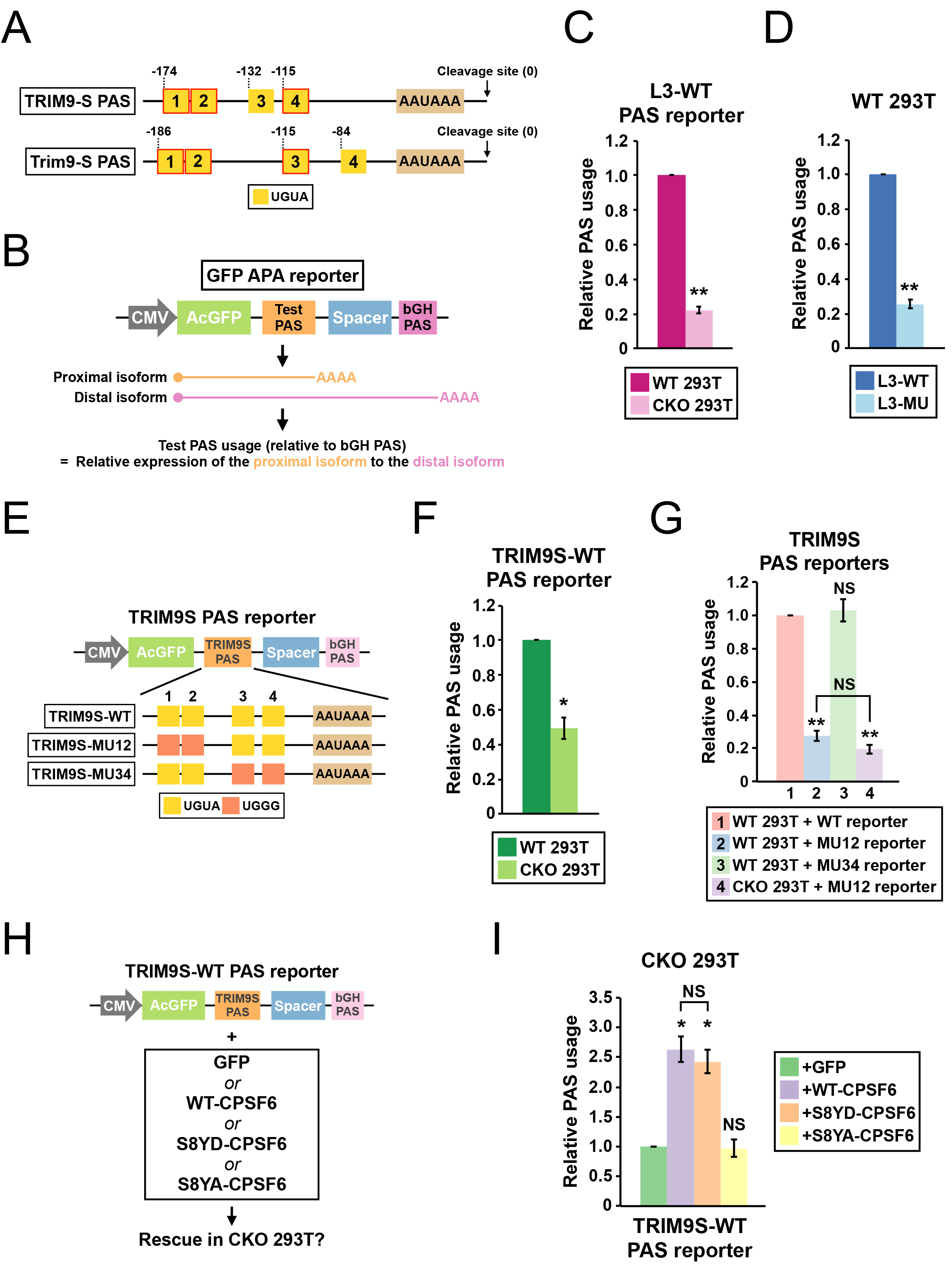
CPSF6 regulates TRIM9-S expression through an evolutionarily conserved twin UGUA motif. (**A**) Illustrations showing human TRIM9-S PAS and mouse Trim9-S PAS, including locations of the UGUA motifs (shown as yellow boxes numbered from 5’ to 3’). The twin UGUA motif is indicated by yellow boxes 1 & 2. The red outline indicates conservation between human and mouse. AAUAAA, the poly(A) signal. See Fig. S4A for the alignment of human and mouse nucleotide sequences. (**B**) Illustrations showing the design of GFP APA reporter to measure the strength of the inserted PAS (Test PAS) by RT-qPCR. See STAR Methods for details. bGH, bovine growth hormone. (**C**) Bar graphs showing usage of L3 wildtype PAS in both wildtype (WT) and CPSF6-KO (CKO) 293T cells from 3 independent experiments (n=3). (**D**) Bar graphs showing usage of both wildtype (L3-WT) and mutant (L3-MU) L3 PASs in 293T cells from 3 independent experiments (n=3). (**E**) Illustrations showing the design of different TRIM9S PAS reporters. (**F**) Bar graphs showing usage of TRIM9-S wildtype PAS in both WT and CKO 293T cells from 3 independent experiments (n=3). (**G**) Bar graphs showing usage of different TRIM9-S PAS reporters in 293T or CKO cells from 3 independent experiments (n=3). (**H**) Illustrations showing the design of rescue experiments using the TRIM9S-WT PAS reporter in CKO cells. (**I**) Bar graphs showing usage of TRIM9S-WT PAS in CKO cells co-transfected with different expression constructs from 3 independent experiments (n=3). All measurements for the PAS reporter assays were performed 24 h after transfection. Error bars indicate SEM. Statistical significance is determined by two-tailed t-test. NS: not significant, *: p<0.05, **: p<0.01.

We first validated the assay system using the L3 PAS, a direct CFIm target with two UGUA motifs 5’ to the poly(A) signal that are crucial for its usage (Zhu et al., 2018). We generated a wildtype L3 PAS reporter (L3-WT)(Fig. S4B). To examine the *trans*-acting effects from CPSF6, we compared L3-WT usage in between wildtype and *CRSF6*-knockout 293T cells (Fig. S4C, referred to as CKO cells thereafter)(Sowd et al., 2016). As expected, usage of L3-WT PAS was strongly decreased in CKO cells (Fig. 4C). Next, to examine the *cis*-acting effects, we generated a mutant L3 PAS reporter (L3-MU) by mutating both UGUAs to UGGGs, a motif previously shown to abolish CFIm regulation (Zhu et al., 2018)(Fig. S4B). Consistent with the previous report (Zhu et al., 2018), usage of L3-MU was much lower compared to L3-WT in 293T cells (Fig. 4D). Notably, the similar levels of decrease in L3 PAS usage between CPSF6 ablation (78%, Fig. 4C) and mutations in both UGUA motifs (75%, Fig. 4D) are consistent with the previous report that these two upstream UGUA motifs in L3 PAS are the main interaction sites with CFIm (Zhu et al., 2018). Altogether, these results demonstrated the sensitivity and validity of our tandem PAS reporter assay in detecting both *cis*- and *trans*-acting effects on PAS regulation.

Next, we proceeded to clone the human TRIM9-S PAS into our reporter (Fig. 4E) and confirmed that the expected cleavage site was used by 3’ RACE (Fig. S4D). To examine the contribution of UGUA motifs to PAS usage, we performed site-directed mutagenesis to generate a series of TRIM9-S PAS reporters with UGGG mutations introduced to different combinations of UGUA motifs (Fig. 4E). Next, we performed reporter assays using all 3 TRIM9-S PAS reporters with the following results: 1) Loss of CPSF6 approximately halved TRIM9-S PAS usage (Fig. 4F, 51% decrease). 2) Mutations in the twin UGUA motif strongly decreased TRIM9-S PAS usage (Fig. 4G, group 1 vs. group 2, 73% decrease). 3) Mutations in the other two UGUA motifs together had no effects on TRIM9-S PAS usage (Fig. 4G, group 1 vs. group 3). 4) Loss of CPSF6 did not statistically significantly change the usage of TRIM9-S PAS with mutated twin UGUA motif (Fig. 4G, group 2 vs. group 4, p=0.12). Altogether, these results support that CPSF6 promotes TRIM9-S PAS usage mostly through the evolutionarily conserved twin UGUA motif.

Phosphorylation of CPSF6 is essential for APA of autophagy genes in *Drosophila* (Fig. S3)(Tang et al., 2018). Therefore, we sought to examine whether phosphorylation of CPSF6 is also required for TRIM9-S PAS usage. We took advantage of the finding that CPSF6 ablation halved TRIM9-S PAS usage (Fig. 4F) and performed genetic rescue experiments in CKO cells. We first obtained WT-CPSF6, S8YD-CPSF6 (a phosphomimetic mutant), and S8YA-CPSF6 (a phosphodeficient mutant) expression plasmids (Jang et al., 2019), and we confirmed that they all expressed similar amount of CPSF6 in 293T cells by western blots (Fig. S4E). We next cotransfected CKO cells with the TRIM9-S WT PAS reporter in addition to each of the 4 following expression plasmids (Fig. 4H): GFP (no rescue control), WT-CPSF6, S8YD-CPSF6, and S8YA-CPSF6. Reporter assays showed that WT-CPSF6 and S8YD-CPSF6, but not S8YA-CPSF6, rescued the loss of TRIM9-S PAS usage (Fig. 4I). These results are consistent with the previous report that CPSF6 is phosphorylated *in vivo* (Zhu et al., 2018) and suggest that phosphorylation of CPSF6 is indeed crucial for its role in promoting TRIM9-S PAS usage. A summary of our characterization of the *mTORC1-CPSF6-Trim9/TRIM9* regulation is provided in Fig. S4F.

### CPSF6-dependent PASs in *BMPR1B* and *MOB4* have functional twin UGUA motifs

Having identified a role of the TRIM9-S twin UGUA motif in the regulation of TRIM9-S PAS by CPSF6, we next asked whether we could find more CPSF6-regulated PASs that contain similar twin UGUA motifs in human. We generated BE2C cells that stably express an shRNA targeting CPSF6 from a tetracycline-inducible promoter (referred to as shCPSF6-BE2C cells thereafter). In shCPSF6-BE2C cells, 72-hour doxycycline treatment strongly reduced CPSF6 mRNA and protein expression as expected (Fig. 5A and 5B). Next, we performed APA profiling using PAPERCLIP (Hwang et al., 2016) on shCPSF6-BE2C cells grown in the absence (high CPSF6) or presence (low CPSF6) of doxycycline treatment to identify CPSF6-dependent PASs (Fig. 5C)(Table S2). We searched for 2-PAS genes that satisfied the following criteria: 1) Switched the major PAS from loss of CPSF6; 2) Had a twin UGUA motif (UGUANUGUA) within 100bp from the cleavage site in the loss-of-use PAS but not the other PAS. We identified 6 candidate genes—all of them have a twin UGUA motif in the distal PAS that lost use with low CPSF6 (Table S3). We chose the top 2 candidates, *BMPR1B* and *MOB4*, for experimental validation (Fig. 5D). *BMPR1B* distal PAS has a 3rd UGUA motif downstream of the UGUAUUGUA motif (Fig. 5E and Fig. S5). In contrast, *MOB4* distal PAS does not have any additional UGUA motif besides the UGUACUGUA motif and a canonical poly(A) signal (Fig. 5E and Fig. S5).

**Figure 5.**
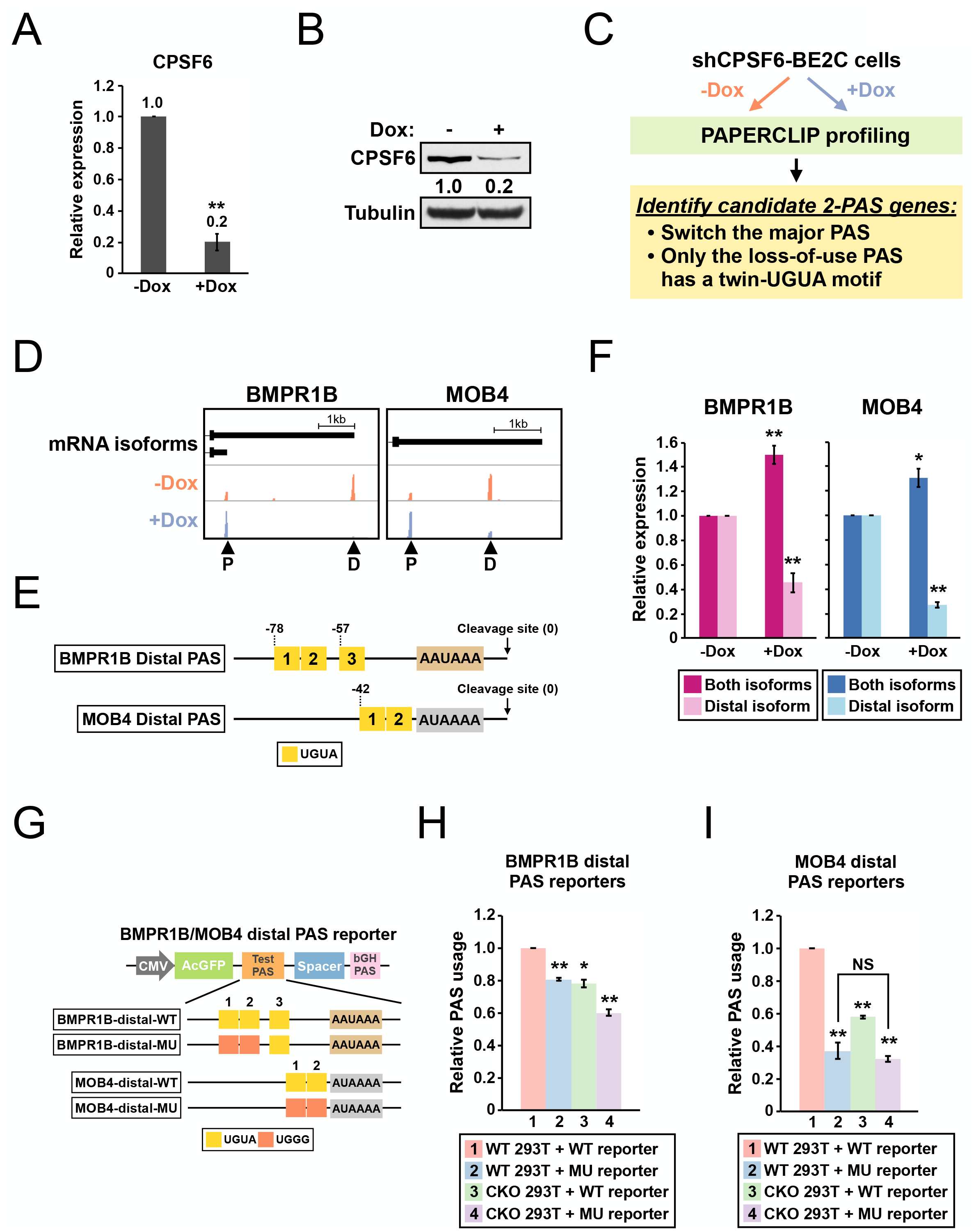
*BMPR1B* and *MOB4* distal PASs are two CPSF6-dependent PASs with a functional twin UGUA motif. (**A**) Bar graphs showing *CPSF6* mRNA expression measured by RT-qPCR in shCPSF6-BE2C cells with (+Dox) and without (-Dox) doxycycline treatment for 72 h from 3 independent experiments (n=3). Dox: 1 μg/mL doxycycline. (**B**) Western blots showing *CPSF6* protein expression in shCPSF6-BE2C cells with and without doxycycline treatment for 72 h. Tubulin: loading control. (**C**) Illustrations showing the experimental strategy to identify candidate genes harboring a functional twin UGUA motif. (**D**) GENCODE annotations and PAPERCLIP results (merged from two biological replicates) in shCPSF6-BE2C cells for *BMPR1B* and *MOB4*. Arrowheads: poly(A) sites identified by PAPERCLIP. P: Proximal. D: Distal. (**E**) Illustrations showing human *BMPR1B* and *MOB4* distal PASs, including locations of the UGUA motifs (shown as yellow boxes numbered from 5’ to 3’). The twin UGUA motif is indicated by yellow boxes 1 & 2. AAUAAA, the poly(A) signal. AUAAAA, a putative non-canonical poly(A) signal. See Fig. S5 for the nucleotide sequences. (**F**) Bar graphs showing expression of *BMPR1B* and *MOB4* mRNA isoforms measured by RT-qPCR in shCPSF6-BE2C cells with (+Dox) and without (-Dox) doxycycline treatment for 72 h from 4 independent experiments (n=4). (**G**) Illustrations showing the design of *BMPR1B* and *MOB4* distal PAS reporters. (**H-I**) Bar graphs showing usage of *BMPR1B* distal PAS (H) or *MOB4* distal PAS (I) in different combinations of 293T cells (WT and CKO) and reporters (WT and MU) 24 h after transfection from 3 independent experiments (n=3). Error bars indicate SEM. Statistical significance is determined by two-tailed t-test. NS: not significant, *: p<0.05, **: p<0.01.

First, we performed RT-qPCR experiments measuring *BMPR1B* and *MOB4* mRNA isoforms in shCPSF6-BE2C cells to verify the loss of distal PAS usage identified by PAPERCLIP. We found that the distal mRNA isoform abundance indeed dropped in doxycycline-treated shCPSF6-BE2C cells for both genes (Fig. 5F). Next, to examine the *cis-*regulatory effects of the identified twin UGUA motifs, we generated *BMPR1B* and *MOB4* PAS reporters that were with or without mutations in the twin UGUA motif (Fig. 5G) for reporter assays in both 293T and CKO cells. The reporter assays showed that: 1) loss of CPSF6 reduced usage of both *BMPR1B* and *MOB4* distal PASs (Fig. 5H and 5I, group 1 vs. group 3, *BMPR1B:* 22% decrease, *MOB4:* 42% decrease); 2) mutations in the twin UGUA motif diminished usage of both *BMPR1B* and *MOB4* distal PASs (Fig. 5H and 5I, group 1 vs. group 2, *BMPR1B:* 19% decrease; *MOB4:* 63% decrease). These results confirm that CPSF6 promotes usage of both *BMPR1B* and *MOB4* distal PASs and demonstrate a functional role of the twin UGUA motif in both PASs. Importantly, loss of CPSF6 did not statistically significantly reduce the usage of *MOB4* distal PAS with the mutant twin UGUA motif (Fig. 5I, group 2 vs. group 4, p=0.45), suggesting that regulation of *MOB4* distal PAS by CPSF6 occurs mostly through the twin UGUA motif just like in TRIM9-S PAS.

### CPSF6 promotes expression of BRD4-L, which has a functional twin UGUA motif in its PAS

The bromodomain protein BRD4 has two major isoforms, BRD4-L and BRD4-S, with opposing biological functions in MDA-MB-231 breast cancer cells (Wu et al., 2020). Although loss of CPSF6 in shCPSF6-BE2C cells did not cause a switch in the major PAS of *BRD4* (which remained to be BRD4-S), our PAPERCLIP profiling did identify BRD4-L PAS as one of the CPSF6-dependent PASs (Fig. 6A). Moreover, although BRD4-L PAS is not conserved in mouse, we found that the sequence arrangement of BRD4-L PAS was similar to that of TRIM9-S PAS— both PASs have a 5’ twin UGUA motif (UGUACUGUA) followed by two individual UGUA motifs at comparable downstream locations (Fig. 6B and Fig. S6A). Therefore, we sought to examine whether CPSF6 also regulates the balance between the two BRD4 isoforms. We first measured BRD4-L and BRD4-S expression in 293T and CKO cells by RT-qPCR and western blots. In 3 independent experiments, we found that the BRD4L-to-BRD4S ratio was consistently lower in CKO cells when compared to 293T cells at both mRNA and protein levels (Fig. 6C-6E), suggesting that CPSF6 promotes BRD4-L expression.

**Figure 6.**
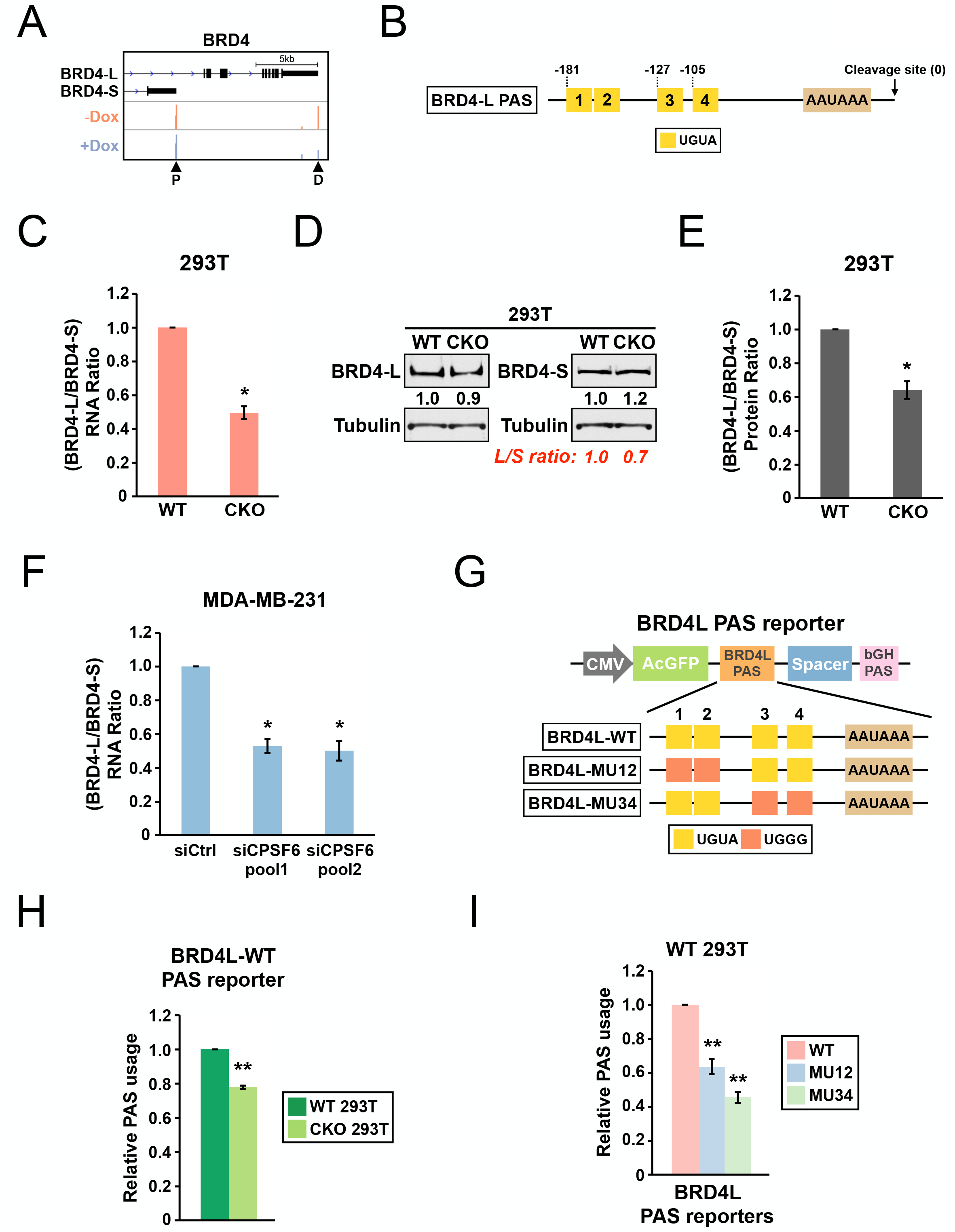
CPSF6 promotes expression of BRD4-L, which has a functional twin UGUA motif in the PAS. (**A**) GENCODE annotations and PAPERCLIP results (merged from two biological replicates) in shCPSF6-BE2C cells for *BRD4*. Arrowheads: poly(A) sites identified by PAPERCLIP. P: Proximal. D: Distal. (**B**) Illustrations showing human BRD4-L PAS, including locations of the UGUA motifs (shown as yellow boxes numbered from 5’ to 3’). The twin UGUA motif is indicated by yellow boxes 1 & 2. AAUAAA, the poly(A) signal. See Fig. S6B for the nucleotide sequence. (**C**) Bar graphs showing the ratio of *BRD4* mRNA isoforms measured by RT-qPCR in 293T and CKO cells from 3 independent experiments (n=3). (**D**) Western blots showing expression of *BRD4* protein isoforms in 293T and CKO cells. Tubulin: loading control. (**E**) Bar graphs showing the ratio of *BRD4* protein isoforms in 293T and CKO cells from 3 independent experiments (n=3). (**F**) Bar graphs showing the ratio of *BRD4* mRNA isoforms measured by RT-qPCR in MDA-MB-231 cells transfected with different siRNAs for 72 h from 3 independent experiments (n=3). siCtrl: control siRNA. (**G**) Illustrations showing the design of BRD4-L PAS reporters. (**H**) Bar graphs showing usage of BRD4-L wildtype PAS in both 293T and CKO cells 24 h after transfection from 4 independent experiments (n=4). (**I**) Bar graphs showing usage of different BRD4-L PAS reporters in 293T cells 24 h after transfection from 4 independent experiments (n=4). Error bars indicate SEM. Statistical significance is determined by two-tailed t-test. *: p<0.05, **: p<0.01.

We next examined whether CPSF6 also regulates BRD4 isoform expression in MDA-MB-231 cells. We acutely knocked down CPSF6 in MDA-MB-231 cells with two different pools of CPSF6-targeting siRNAs (Fig. S6B). Similar to the 293T/CKO experiments, we observed a consistent decrease in the BRD4L-to-BRD4S mRNA ratio in MDA-MB-231 cells from 3 independent experiments (Fig. 6F). Moreover, the magnitude of decrease in the BRD4L-to-BRD4S mRNA ratio from loss of CPSF6 was similar between 293T cells and MDA-MB-231 cells (Fig. 6C and Fig. 6F). To facilitate western blot analysis on BRD4 protein isoform expression, we generated shCPSF6-MDA-MB-231 cells that stably express an shRNA targeting CPSF6 from a tetracycline-inducible promoter (same as shCPSF6-BE2C cells). As expected, in shCPSF6-MDA-MB-231 cells, doxycycline treatment strongly suppressed CPSF6 mRNA expression and again reduced the mRNA ratio of BRD4-L over BRD4-S (Fig. S6C). We next proceeded to examine BRD4-L and BRD4-S protein expression in shCPSF6-MDA-MB-231 cells with and without doxycycline treatment. Surprisingly, in contrast to the results from 293T/CKO cells, we did not find consistent expression changes between BRD4-L and BRD4-S proteins in response to doxycycline treatment (data not shown). Nevertheless, these results from MDA-MB-231 and shCPSF6-MDA-MB-231 cells provide additional evidence to support that CPSF6 promotes BRD4-L mRNA expression.

Next, as we previously did with TRIM9-S PAS (Fig. 4E), we generated a series of BRD4-L PAS reporters to examine the contribution of UGUA motifs to BRD4-L PAS usage (Fig. 6G). Consistent with the results from PAPERCLIP profiling, the WT BRD4-L PAS showed a consistent decrease in usage in CKO cells when compared to 293T cells (Fig. 6H, 22% decrease). Furthermore, similar to TRIM9-S PAS, mutations in the twin UGUA motif indeed reduced BRD4-L PAS usage (Fig. 6I, MU12: 36% decrease), supporting a role of the twin UGUA motif in promoting BRD4-L PAS usage. However, mutations in the two downstream UGUA motifs in BRD4-L PAS also reduced BRD4-L PAS usage (Fig. 6I, MU34: 54% decrease), which was not the case for TRIM9-S PAS (Fig. 4G). We concluded that, like TRIM9-S PAS, BRD4-L PAS is regulated by CPSF6 and has a functional twin UGUA motif. Nevertheless, TRIM9-S PAS and BRD4-L PAS are not regulated by CPSF6 in the same way despite similar UGUA motif arrangements. Importantly, we note that, through PAPERCLIP profiling and reporter assays in *BMPR1B, MOB4* and BRD4-L, we have expanded the number of human CPSF6-dependent PASs harboring a functional twin UGUA motif beyond TRIM9-S.

### Insertion of a twin UGUA motif into the *JUNB* PAS is sufficient to confer regulation by CPSF6 and mTORC1

We next wanted to test whether insertion of the TRIM9-S twin UGUA motif into a heterologous PAS is sufficient to make the host PAS responsive to CPSF6 and mTORC1 regulation in our reporter assay. For the host PAS, we selected human *JUNB* PAS, which does not contain any UGUA motif, and its usage is not affected by loss of CPSF6 (Hwang et al., 2016)(Fig. 7A and Fig. S7A). We found that insertion of the TRIM9-S twin UGUA motif strongly increased the *JUNB* PAS usage (Fig. 7B, group 2, 84% increase) but not the twin UGGG mutant (Fig. 7B, group 5, 37% decrease) in 293T cells. Importantly, this increase was entirely dependent on CPSF6 as it was completely abolished in CKO cells (Fig. 7B, group 3). Interestingly, insertion of a single UGUA motif did not statistically significantly alter the *JUNB* PAS usage (Fig. 7B, group 4, p=0.18), suggesting that a second copy of UGUA is essential for the biological activity of the TRIM9-S twin UGUA motif.

**Figure 7.**
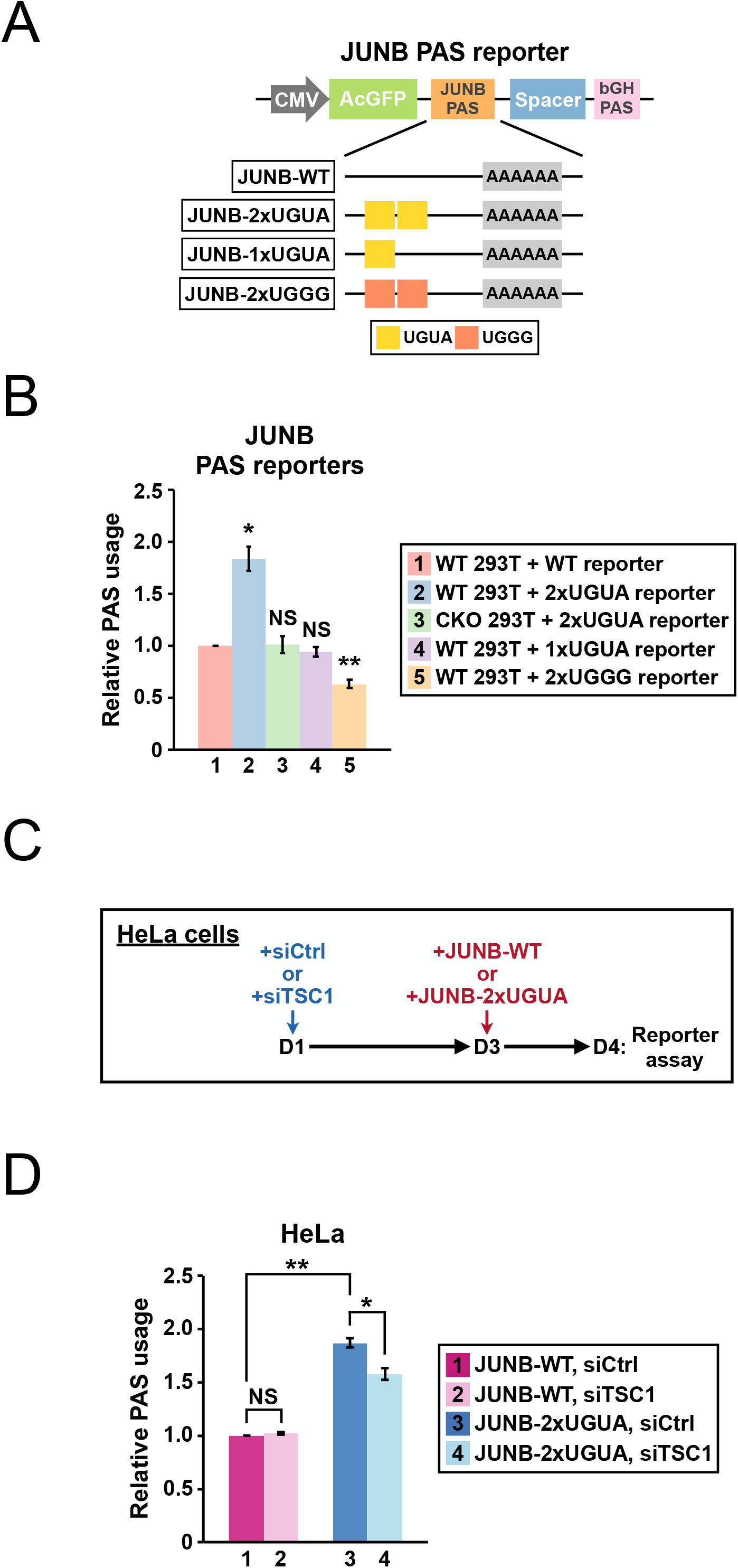
Insertion of a twin UGUA motif into the *JUNB* PAS is sufficient to confer regulation by CPSF6 and mTORC1. (**A**) Illustrations showing the design of *JUNB* PAS reporters. (**B**) Bar graphs showing usage of different *JUNB* PAS reporters (WT, 2xUGUA, 1xUGUA and 2xUGGG) in 293T or CKO cells 24 h after transfection from 3 independent experiments (n=3). (**C**) Illustrations showing the experimental design for the *JUNB* PAS reporter assay in HeLa cells. (**D**) Bar graphs showing usage of JUNB-WT and JUNB-2xUGUA PAS reporters in HeLa cells with normal (siCtrl) or hyperactive (siTSC1) mTORC1 from 3 independent experiments (n=3). Error bars indicate SEM. Statistical significance is determined by two-tailed t-test. NS: not significant, *: p<0.05, **: p<0.01.

Next, we sought to extend the *JUNB* PAS reporter assay to test whether presence of the TRIM9-S twin UGUA motif makes the *JUNB* PAS become sensitive to different levels of mTORC1 activities. We hypothesized that CPSF6 inhibition from mTORC1 hyperactivation would dampen usage of JUNB-2xUGUA PAS but not JUNB-WT PAS. Because HeLa cells offer a better model than 293T cells for mTORC1 hyperactivation (Alesi et al., 2021), we transfected HeLa cells with control siRNAs or *TSC1* siRNAs to hyperactivate mTORC1 48 hours prior to a second transfection of either JUNB-WT or JUNB-2xUGUA reporters (Fig. 7C). *TSC1* knockdown is indistinguishable between the two experimental groups of JUNB-WT and JUNB-2xUGUA (Fig. S7B). As seen in 293T cells (Fig. 7B), usage of JUNB-2xUGUA is strongly increased compared to that of JUNB-WT in HeLa cells transfected with control siRNAs (Fig. 7D, group 1 vs. group 3, 87% increase). While usage of JUNB-WT was not affected by mTORC1 hyperactivation (Fig. 7D, group 1 vs. group 2, p=0.18), usage of JUNB-2xUGUA was consistently decreased in siTSC1-transfected HeLa cells compared to siCtrl-transfected HeLa cells (Fig. 7D, group 4 vs. group 3, 29% decrease). Taken together, these results demonstrate that insertion of the TRIM9-S twin UGUA motif into a heterologous PAS is sufficient to make the host PAS become responsive to CPSF6 expression and mTORC1 activities.

## DISCUSSION

In this study, we successfully applied the powerful cTag-PAPERCLIP technique to a well-established TSC mouse model and identified an evolutionarily conserved regulatory relationship between the mTORC1 pathway and *Trim9/TRIM9*. Our results demonstrate the feasibility of our approach to circumvent the cumbersome breeding process and model constitutive mTORC1 signaling, the hallmark of TSC (Salussolia et al., 2019), in desired cell populations in the mouse brain *in vivo*. Deletion of *Tsc1* in different brain cell types in mouse generates distinct phenotypes, each capturing separate human disease features (Bateup et al., 2013; Benthall et al., 2021; Ercan et al., 2017; Jiang et al., 2016; Kosillo et al., 2019; Lebrun-Julien et al., 2014; Meikle et al., 2007; Tsai et al., 2012; Uhlmann et al., 2002; Zou et al., 2017). Our results suggest that extending APA analysis to additional cell types in TSC brain with the rapidly expanding AAV toolkit (Challis et al., 2022) would facilitate identification of possible molecular defects in TSC due to APA dysregulation in addition to generating a panoramic view of APA regulation in TSC.

We performed extensive characterizations to genetically delineate the pathway linking mTORC1 to *Trim9/TRIM9* isoform regulation all the way down to the level of *cis*-regulatory motif (Fig. S4F). We found similarities between Trim9/TRIM9-S regulation by mTORC1 and the previously reported regulation of *Drosophila* autophagy genes, *Atg1* and *Atg8a* (Fig. S3)(Tang et al., 2018). For example, CDK8 and the phosphorylation status of CPSF6 are also important factors for Trim9/TRIM9-S regulation by mTORC1 (Fig. 3G and 4I). However, we note that the APA shifts in both *Atg1* and *Atg8a* lead to 3’ UTR lengthening and mRNA stabilization while the APA shift in *Trim9/TRIM9* changes gene isoform expression—Trim9/TRIM9-L and Trim9/TRIM9-S proteins have distinct 3’ termini (Fig. 1C) and different biological functions (see below). Therefore, the mTORC1 signaling pathway could reprogram gene expression both qualitatively and quantitatively solely through APA regulation.

It is very intriguing that the regulation of *Trim9/TRIM9* by CPSF6 seems to be unconventional in multiple ways. First, it is widely known that loss of CPSF6 mainly results in proximal APA shifts (Ghosh et al., 2022; Gruber et al., 2012; Hwang et al., 2016; Li et al., 2015; Martin et al., 2012; Masamha et al., 2014; Zhu et al., 2018). For example, in our current (shCPSF6-BE2C cells) and previous studies (HeLa and LN229 cells)(Hwang et al., 2016), more than 90% of APA shifts in 2-PAS genes from CPSF6 knockdown are proximal shifts. However, for *Trim9/TRIM9*, inhibition of CPSF6 from mTORC1 hyperactivation results in a distal APA shift instead of a proximal APA shift (Fig. 1B). Therefore, *Trim9/TRIM9* seems to belong to a small minority of CPSF6 targets in which the main regulation site is the proximal PAS. Second, the *cis*-regulatory motif in the Trim9/TRIM9-S PAS is not the more common UGUA motif but an evolutionarily conserved UGUAYUGUA motif. Our JUNB reporter assay results suggest that it might be advantageous to have a second copy of UGUA, which is necessary for biological effects in this particular context (Fig. 7B). This is consistent with a previous study that the chance for NUDT21 binding is increased by having a second copy of UGUA in the RNA (Yang et al., 2010). The UGUAYUGUA motif is reminiscent of a “bipartite motif”, which is a pair of short motifs spaced one or more bases apart and the type of motif preferred by some RNA-binding proteins with multiple RNA-binding domains (Dominguez et al., 2018; Zhang and Darnell, 2011). Although NUDT21 forms a dimer (each contains a Nudix domain for RNA binding) that can bind to two UGUA motifs simultaneously on the same RNA (Yang et al., 2010), the minimal distance between the two UGUA motifs that permits simultaneous binding is 7 nucleotides (Yang et al., 2011b). Therefore, it seems unlikely that the UGUAYUGUA motif functions by engaging both NUDT21 proteins in the dimer at the same time. Lastly, in human and mouse PASs, the UGUA motif is enriched in the 40~100 bp region upstream the cleavage site with a peak around 50 bp upstream the cleavage site (Gruber and Zavolan, 2019; Wang et al., 2018; Zhu et al., 2018). In contrast, the Trim9/TRIM9-S UGUAYUGUA motif is located more than 170 bp upstream of the cleavage site (Fig. 4A). It will be interesting to understand how CFIm promotes 3’-end processing from this location in the future. It is worth noting that the mRNA 3’-end region is often folded, and forming secondary structures can promote 3’-end processing by shortening the actual distance to the cleavage site (Wu and Bartel, 2017).

We provide multiple lines of evidence to support that the UGUAYUGUA motif is not distinct to *Trim9/TRIM-9*. First, we have identified and experimentally confirmed the existence of a functional UGUAYUGUA motif in additional human CPSF6-dependent PASs (Table S3). Second, despite differences in the motif locations and the surrounding sequence contexts, the UGUAYUGUA motif plays a crucial role in CPSF6-mediated regulation in both *MOB4* distal PAS and TRIM9-S PAS (Fig. 4G and 5I). Lastly, the UGUAYUGUA motif can be transferred to a heterologous PAS to gain CPSF6-mediated regulation (Fig. 7B). Our findings indicate that the UGUAYUGUA motif is a naturally occurring *cis*-regulatory motif for CPSF6 in human and mouse that might be stronger than a single UGUA motif. Furthermore, our results suggest that, in order to fully grasp how CFIm regulates APA, it might be necessary to look beyond the simple UGUA motif and take different combinations of UGUA motifs into consideration as well.

The functional difference between Trim9/TRIM9-L and Trim9/TRIM9-S has been reported in a few different biological contexts including neuron morphogenesis (Liu et al., 2018; Menon et al., 2015; Qin et al., 2016; Winkle et al., 2014). In neurons, Trim9-L protein controls Netrin-mediated axon branching—it suppresses axon branching in the absence of Netrin-1 but permits it when Netrin-1 is present (Winkle et al., 2014). The response of Trim9-L to Netrin-1 depends on the Trim9-L SPRY domain, which interacts with the Netrin-1 receptor, DCC (Winkle et al., 2014). Because the Trim9-S protein lacks the SPRY domain, in *Trim9*-KO neurons rescued with Trim9-S protein, the axon branching is constantly suppressed and unresponsive to Netrin-1 (Menon et al., 2015). Interestingly, in the mouse embryonic brain, Trim9-S protein seems to be the predominant isoform (Winkle et al., 2014; 2016). We speculate that a premature increase in Trim9-L protein due to mTORC1 hyperactivation in the susceptible developmental window might alter normal neuron morphogenesis and synapse formation. More studies will be needed to elucidate the functional significance of *Trim9/TRIM9* isoform imbalance in TSC.

Taken together, our study reveals that mTORC1 hyperactivation causes *Trim9/TRIM9* isoform imbalance in TSC. Furthermore, we show that the UGUAYUGUA motif in the Trim9/TRIM9-S PAS and CFIm are key *cis*- and *trans*-acting factors linking mTORC1 to the balance between *Trim9/TRIM9* isoforms, respectively. Importantly, although transcriptional and translational abnormalities have been identified in *Tsc1-* and *Tsc2*-deficient neurons (Dalal et al., 2021; Nie et al., 2015), our results demonstrate that gene isoform imbalance is another mechanism for hyperactive mTORC1 to alter normal gene expression. As widespread dysregulation of gene isoform expression was recently found in cancer and neurologic disorders (Gandal et al., 2018; Kahles et al., 2018), our results suggest that investigation of possible gene isoform imbalance in mTORopathies and cancer with hyperactive mTORC1 might provide important biological insights.

## Acknowledgements

We would like to thank Dr. Alan Engelman for providing CPSF6 expression plasmids and CKO HEK293T cells. We thank the Division of Laboratory Animal Resources (DLAR) at the University of Pittsburgh for technical assistance in mouse injection and the Vector Core at the University of Pennsylvania for AAV production. We acknowledge the Health Sciences Sequencing Core at UPMC Children’s Hospital of Pittsburgh the University of Pittsburgh Center for Research Computing, and the University of Pittsburgh HSCRF Genomics Research Core for high-throughput sequencing and Sanger sequencing. This work was supported by a grant from the National Institutes of Health (NS113861 to H-W.H.).

## Author Contributions

Conceptualization, H.-W.H.; Methodology, R.S.H. and H.-W.H.; Formal analysis, R.S.H. and H.-W.H.; Investigation, R.S.H., A.K.K., J.R.M., V.I.A., and H.-W.H.; Writing—original draft preparation: H.-W.H.; Writing—review and editing: R.S.H. and H.-W.H.; Supervision: H.-W.H.

## METHODS

### Cell culture

N2a, HEK293T, CKO HEK293T, MDA-MB-231, and HeLa cells were grown in Dulbecco’s modified Eagle’s medium (DMEM). BE2C cells were grown in DMEM/F12. All media were supplemented with 10% FBS and penicillin-streptomycin. CKO HEK293T cells were provided by Dr. Alan Engelman. shCPSF6-BE2C and shCPSF6-MDA-MB-231 cells were generated by transducing wildtype BE2C and MDA-MB-231 cells with lentiviruses encoding a short hairpin RNA targeting CPSF6 (EZ-Tet-shCPSF6-Puro, see Method Details below) followed by puromycin selection. Cpsf6-KD N2a cells were generated by transfecting N2a cells that stably express Cas9 and a Cre-inducible blasticidin cassette with a plasmid encoding Cre and sgRNAs targeting Cpsf6 followed by double selection with blasticidin and puromycin. siRNA transfection was performed using DharmaFECT reagents (Horizon Discovery) with Silencer Select siRNAs (Invitrogen) or ON-TARGETplus siRNAs (Horizon Discovery) at the final concentration of 10 or 25 nM following manufacturer’s instructions. All siRNAs used are listed in Table S5. Plasmid transfection was performed using X-tremeGENE9 (MilliporeSigma) following manufacturer’s instructions.

Doxycycline was obtained from Sigma and was used at 1 μg/mL. Senexin A was obtained from Tocris Bioscience and was used at 50μM. Torin 1 and TG003 were obtained from Cayman Chemical and were used at 250nM (Torin 1) or 50μM (TG003).

### Mouse

All procedures were conducted according to the Institutional Animal Care and Use Committee (IACUC) guidelines at the University of Pittsburgh. Camk2a-Cre and *Tsc1*-floxed mice (Tsc1^tm1Djk/J^) were obtained from the Jackson Lab and were maintained as homozygotes. cTag-PABP mice were obtained from the Rockefeller University and were maintained by backcrossing to C57BL/6J. For cTag-PAPERCLIP profiling, adult (8-12 week-old) *Tsc1*-wildtype or *Tsc1*-floxed cTag-PABP mice of both sexes received one-time retro-orbital injection with AAVs expressing iCre from mouse *Camk2a* promoter (pAAV-Camk2a-iCre) at the dose of 1X10^12^ genome copies (gc) unless specified, and they were housed for 2~3 weeks before sacrifice. AAV was generated and packaged with the PHP.eB capsid by the University of Pennsylvania Vector Core. For *Tsc1*-floxed cTag-PABP mice, successful activation of mTORC1 signaling from injection was verified by S6 and Phosphor-S6 western blots before cTag-PAPERCLIP profiling. For Fig. S1E, *Camk2a-Cre; Tsc1^fl/fl^* mice were sacrificed at 4 weeks of age because their survival decreased sharply afterwards (Bateup et al., 2013).

### cTag-PAPERCLIP, PAPERCLIP and informatics analysis

cTag-PAPERCLIP and PAPERCLIP library construction was performed as previously described (Hwang et al., 2017; Kunisky et al., 2021). For cTag-PAPERCLIP, mouse brain cortices were crosslinked with 254 nm UV at 400 mJ/cm^2^, lysed in 1X PXL Buffer (1X PBS, 0.1% SDS, 0.5% NP-40, 0.5% Sodium deoxycholate). For PAPERCLIP, cultured cells were crosslinked with 254 nm UV at 200 mJ/cm^2^, and lysed in 1X TS Buffer (1X PBS, 0.1% SDS, 1.0% Triton X-100). Both mouse brain and cell lysates were digested with DNase I (Promega) for 5 minutes at 37°C and then with RNase A (ThermoFisher) for 5 minutes at 37°C. Lysates were cleared by centrifugation at 20,000 x g at 4°C for 10 min.

For cTag-PAPERCLIP, PABP-GFP-mRNA complexes were immunoprecipitated from cleared lysates for 2 hours at 4°C using Dynabeads protein G (ThermoFisher) conjugated to anti-GFP (clones 19F7 and 19C8, MSKCC). Beads were then washed sequentially with 1X PXL Buffer, 5X PXL Buffer (5X PBS, 0.1% SDS, 0.5% NP-40, 0.5% Sodium deoxycholate) and PNK buffer (50mM Tris-HCl, pH 7.4, 10mM MgCl_2_, 0.5% NP-40). For PAPERCLIP, endogenous PABP-mRNA complexes were immunoprecipitated from cleared lysates for 2 hours at 4°C using Dynabeads protein G (ThermoFisher) conjugated to anti-PABP (clone 10E10, Sigma). Beads were then washed sequentially with 1X TS Buffer, 2X TS Buffer (2X PBS, 0.1% SDS, 1.0% Triton X-100) and PNK buffer (50mM Tris-HCl, pH 7.4, 10mM MgCl_2_, 0.5% NP-40). After wash, for both cTag-PAPERCLIP and PAPERCLIP, the immunoprecipitated protein-RNA complexes were treated with alkaline phosphatase and 5’ labeled with ^32^P-gamma-ATP using T4 Polynucleotide Kinase on beads. The protein-RNA complexes were then eluted from beads, resolved on a Bis-Tris NuPAGE gel (8% for cTag-PAPERCLIP; 10% for PAPERCLIP), transferred to a nitrocellulose membrane, and film-imaged. Regions of interest were excised from the membrane and the RNA was isolated by Proteinase K digestion and phenol/chloroform extraction. Eluted RNA was reverse-transcribed using SuperScript III with BrdUTP. The resulting cDNAs were purified by two rounds of immunoprecipitation with Dynabeads protein G conjugated to anti-BrdU (clone IIB5, Millipore). The purified cDNAs were then ligated using CircLigase II (Lucigen) and PCR-amplified to generate the sequencing library.

Individual cTag-PAPERCLIP or PAPERCLIP libraries were multiplexed and sequenced by NextSeq (Illumina) to obtain 125-nt single-end reads. The procedures for raw read processing, mapping and poly(A) site annotation were previously described (Hwang et al., 2017). The raw reads were processed (filtered and collapsed) using the CIMS package. Poly(A) sequence at the 3’ end was trimmed using CutAdapt. Trimmed reads that are longer than 25 nucleotides are aligned to mouse (mm10) or human (hg19) genome using Novoalign. The aligned reads were further processed using the CIMS package to remove PCR duplicates and to cluster overlapping reads for poly(A) site identification. For APA shift analysis, different cutoffs for two poly(A) site genes were computed to generate a broad set of two poly(A) site genes for comparison. Significant APA shift is defined as previously described (Hwang et al., 2016): FDR < 0.05 and a greater than 2-fold change of (proximal PAS/distal PAS) ratio between experimental conditions. All gene lists are provided (Table S1 and S2). Kallisto (Bray et al., 2016) was used to estimate TRIM9 isoform abundance in GSE78961 (Fig. 1F).

### SDS-PAGE and western blots

20~60μg lysates from culture cells or mouse tissues were separated on 10% Bis-Tris or 3~8% Tris-Acetate Novex NuPAGE gels (Invitrogen) and transferred to nitrocellulose membrane following standard procedures. The following antibodies are used for western blotting: mouse monoclonal anti-beta actin (Proteintech, 66009-1-Ig), mouse monoclonal anti-alpha tubulin (Millipore, CP06), mouse monoclonal anti-HA (clone 16B12, BioLegend, 901501), rabbit polyclonal anti-TRIM9 (Proteintech, 10786-1-AP), rabbit monoclonal anti-S6 ribosomal protein (Cell Signaling Technology, 2217), rabbit monoclonal anti-Phospho-S6 ribosomal protein (Ser240/244) (Cell Signaling Technology, 5364), rabbit polyclonal anti-CPSF6 (Bethyl Labs, A301-356A), rabbit polyclonal anti-BRD4 (Bethyl Labs, A301-985A), rabbit monoclonal anti-BRD4 (Abcam, ab128874).

### Reverse transcription and quantitative PCR (RT-qPCR)

qPCR was performed using PerfeCTa SYBR Green SuperMix (QuantaBio) in triplicates. All primer sequences are listed in Table S4. For mRNA quantification, reverse transcription was performed using ProtoScript II First Strand cDNA Synthesis Kit (NEB) using d(T)_23_VN primer with DNase I (Invitrogen) digestion on 1 μg total RNA generated from Trizol (Invitrogen) extraction. The cycling parameters for qPCR were: 95°C for 10 min. followed by 40 cycles of 95°C for 15 sec., 58°C for 30 sec., 72°C for 20 sec. Quantification was calculated using the ΔΔCt method with the following endogenous controls: ACTB (human) and Rplp0 (mouse).

For PAS reporter assay, reverse transcription was performed using ProtoScript II First Strand cDNA Synthesis Kit (New England Biolabs) using an anchored d(T) primer (R1-T25-VN) with DNase I (Invitrogen) digestion on 1 μg total RNA generated from Trizol (Invitrogen) extraction. qPCR quantification of the proximal and distal mRNA isoforms generated from each PAS reporter is performed using a PAS specific forward primer and a common reverse primer complementary to the anchoring sequence of R1-T25-VN. The cycling parameters for qPCR were: 95°C for 10 min. followed by 40 cycles of 95°C for 15 sec., 58°C for 30 sec., 72°C for 8 sec. Quantification was calculated using the ΔΔCt method relative to the distal mRNA isoform that uses the bGH PAS.

### Cloning and constructs

Standard cloning procedure (restriction digest, ligation and transformation) was performed to generate desired constructs. All insert sequences were verified by Sanger sequencing. Oligonucleotides and primers are listed in Table S4. pIRES2-EGFP-CPSF6[551], pIRES2-EGFP-CPSF6[S8YA], pIRES2-EGFP-CPSF6[S8YD] were provided by Dr. Alan Engelman. pAAV-Camk2a-iCre was generated by replacing GFP in pAAV-CAMKII-GFP (Addgene, 64545) with iCre (a codon-optimized Cre recombinase), which was amplified from pEMS1985 (Addgene, 49116) by PCR. EZ-Tet-shCPSF6-Puro was generated by inserting a short hairpin RNA targeting CPSF6 into EZ-Tet-pLKO-Puro (Addgene, 85966) between the NheI and EcoRI sites. GFP APA reporter was generated through the following modifications of the pcDNA 3.1 plasmid: 1) insertion of AcGFP between the NheI and XbaI sites, and 2) insertion of a spacer sequence in front of the bGH poly(A) signal using the BamHI site. The following PAS reporters were generated by insertion of oligonucleotides into the GFP APA reporter between the XhoI and XbaI sites: L3-WT, L3-MU, BMPR1B-distal-WT, BMPR1B-distal-MU, MOB4-distal-WT, MOB4-distal-MU, JUNB-WT, JUNB-2xUGUA, JUNB-1xUGUA, JUNB-2xUGGG. TRIM9S-WT and BRD4L-WT PAS reporters were generated by inserting TRIM9-S and BRD4-L PASs (amplified from human genomic DNA) into the GFP APA reporter between the XhoI and XbaI sites. TRIM9S-MU12, TRIM9S-MU34, BRD4L-MU12, and BRD4L-MU34 PAS reporters were generated by site-directed mutagenesis of TRIM9S-WT or BRD4L-WT PAS reporters using Q5 Site-Directed Mutagenesis Kit (New England Biolabs).

## QUANTIFICATION AND STATISTICAL ANALYSIS

Details of statistical tests are indicated below and in the Figure Legends. Statistical analyses were performed using R.

For Figures 1F (left panel), 2B, 2D, 2F, 2G, 3B-G, 4C-D, 4F-G, 4I, 5A, 5F, 5H-I, 6C, 6E-F, 6H-I, 7B, 7D, S1C, S2B-C, S6B-C, S7B, statistical significance is determined by two-tailed Welch Two Sample t-test.

For Figures 1F (right panel), statistical significance is determined by one-tailed Welch Two Sample t-test.

For all figures: *: p < 0.05, **: p < 0.01.

## SUPPLEMENTARY FIGURE LEGENDS

**Figure S1.**
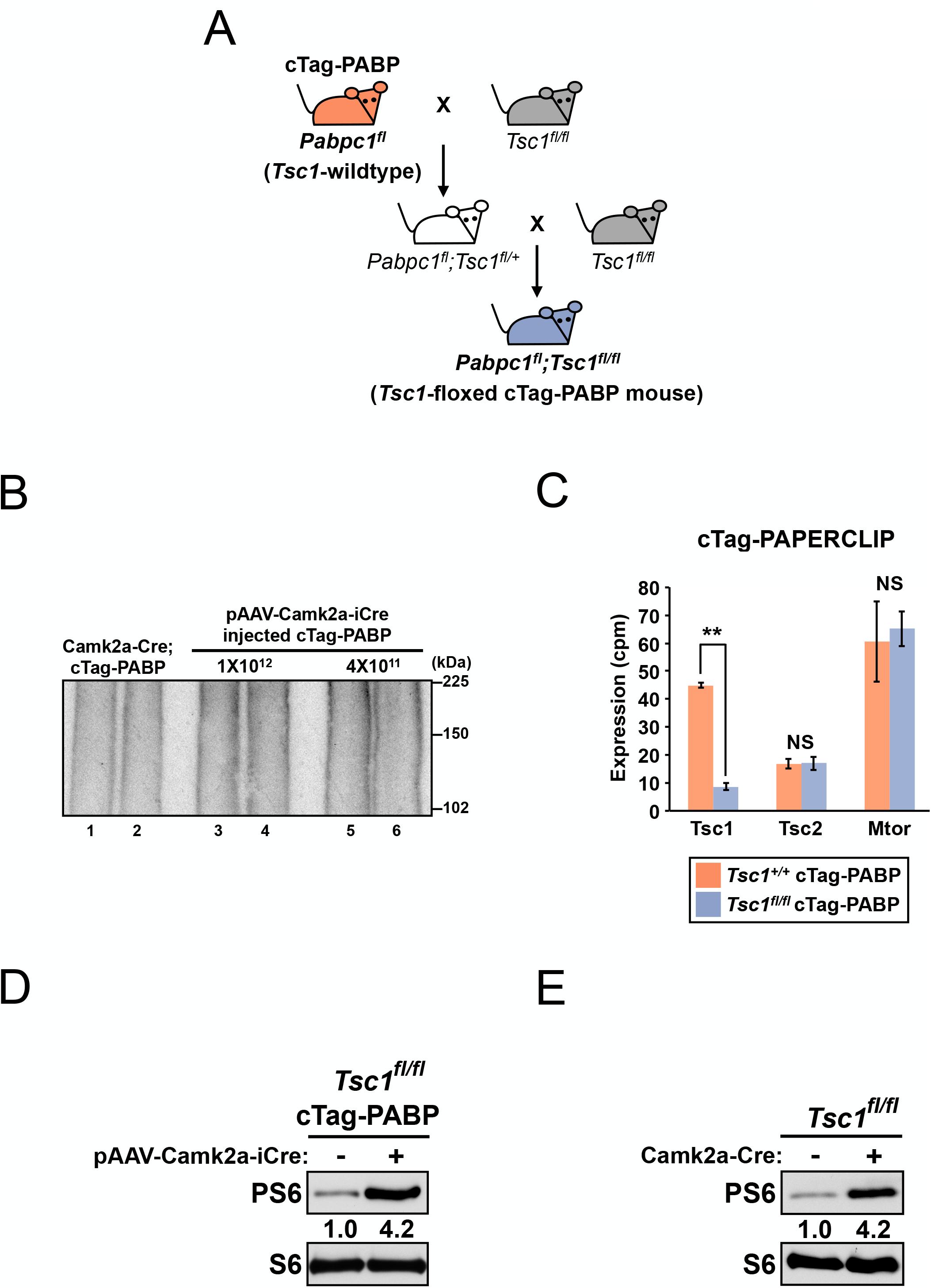
Systemic AAV delivery of Cre recombinase activates PABP-GFP expression and mTORC1 signaling with similar efficiency to genetic breeding, related to Figure 1. (**A**) The breeding strategy to generate *Tsc1*-floxed cTag-PABP mice. (**B**) Autoradiography from a cTag-PAPERCLIP experiment using brain cortices from: 1) a *Camk2a-Cre;* cTag-PABP mouse (lanes 1 & 2), 2) a cTag-PABP mouse injected with pAAV-Camk2a-iCre (1×10^12^ gc)(lanes 3 & 4), and 3) a cTag-PABP mouse injected with pAAV-Camk2a-iCre (4×10^11^ gc)(lanes 5 & 6). gc, genome copies. (**C**) Bar graphs showing quantitation of *Tsc1, Tsc2* and *Mtor* mRNA expression from two biological replicate cTag-PAPERCLIP experiments in pAAV-Camk2a-iCre injected cTag-PABP mice (*Tsc1*-wildtype or *Tsc1*-floxed). The color scheme is the same as in (A). (**D**) Western blots showing expression of total and phosphorylated S6 (PS6) ribosomal protein in the brain cortices of adult *Tsc1*-floxed cTag-PABP mice (with or without pAAV-Camk2a-iCre injection). (**E**) Western blots showing expression of total and phosphorylated S6 ribosomal protein in the brain cortices of a 4-week-old *Camk2a-Cre; Tsc1*-floxed mouse and a *Tsc1*-floxed littermate. Error bars indicate SEM. Statistical significance is determined by two-tailed t-test. NS: not significant, **: p<0.01.

**Figure S2.**
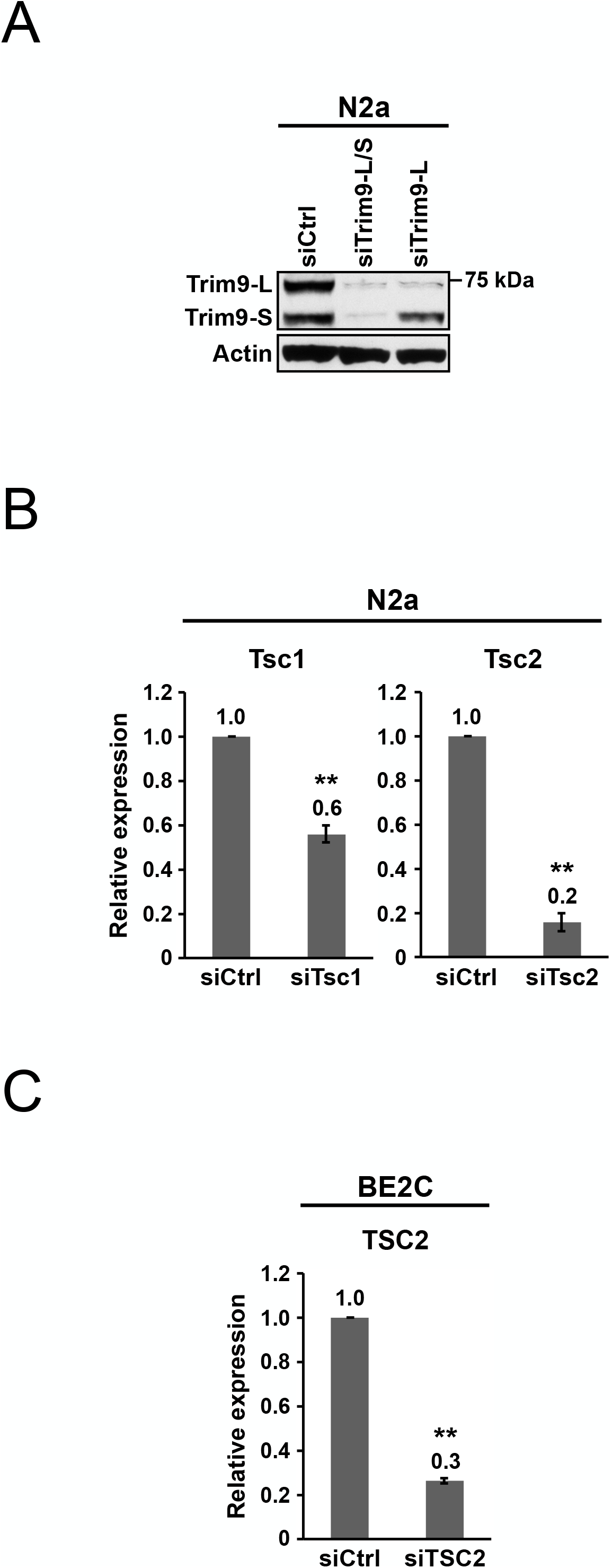
Characterization of the TRIM9 antibody and measurements of *Tsc1/Tsc2* siRNA knockdown efficiency, related to Figure 2. (**A**) Western blots showing expression of Trim9-L and Trim9-S proteins in N2a cells transfected with different siRNAs for 72 h. siCtrl: control siRNA. Actin: loading control. (**B**) Bar graphs showing the knockdown efficiency of *Tsc1* and *Tsc2* siRNAs measured 72 h after transfection by RT-qPCR in N2a cells in the same experiments shown in Fig. 2F (n=4). (**C**) Bar graphs showing the knockdown efficiency of *TSC2* siRNAs measured 72 h after transfection by RT-qPCR in BE2C cells in the same experiments shown in Fig. 2G (n=4). Error bars indicate SEM. Statistical significance is determined by twotailed t-test. **: p<0.01.

**Figure S3.**
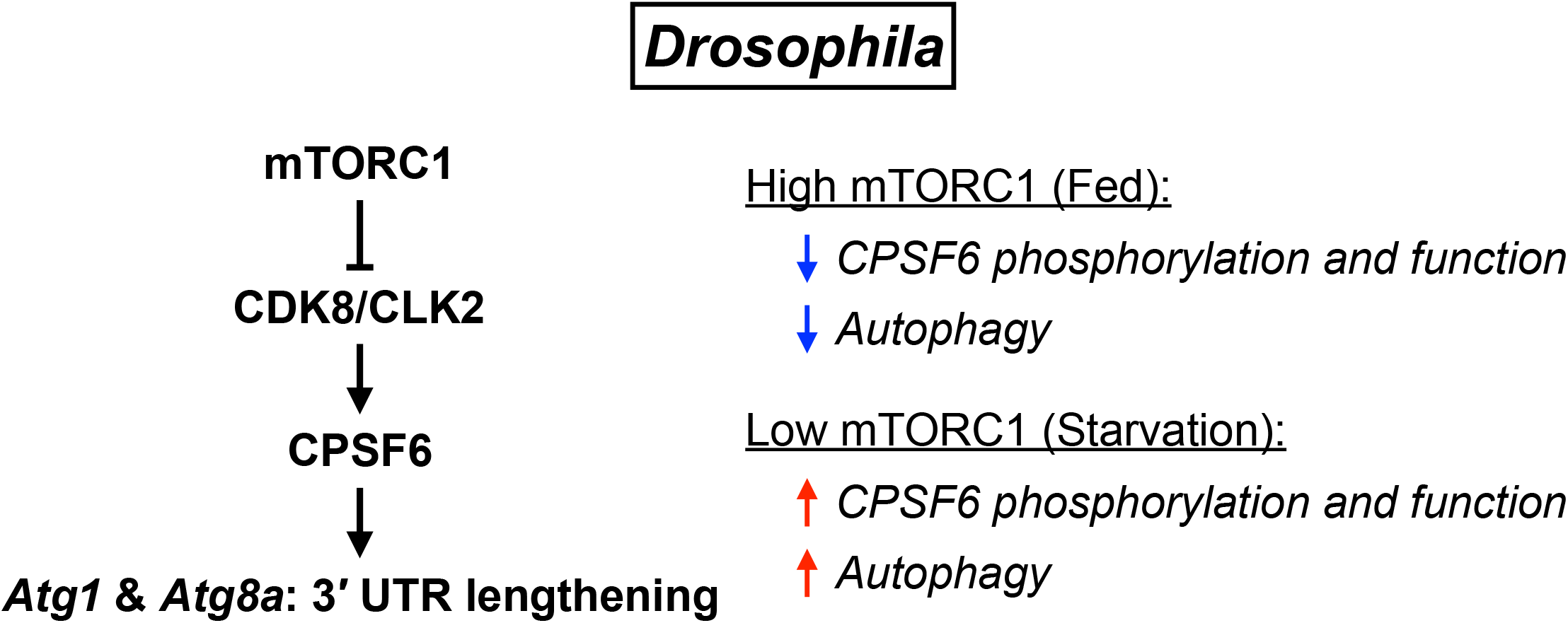
Reported mTORC1 regulation on autophagy genes through CPSF6 in *Drosophila*, related to Figure 3. Illustrations showing the mTORC1 regulation on APA of autophagy genes as reported in Tang et al. 2018.

**Figure S4.**
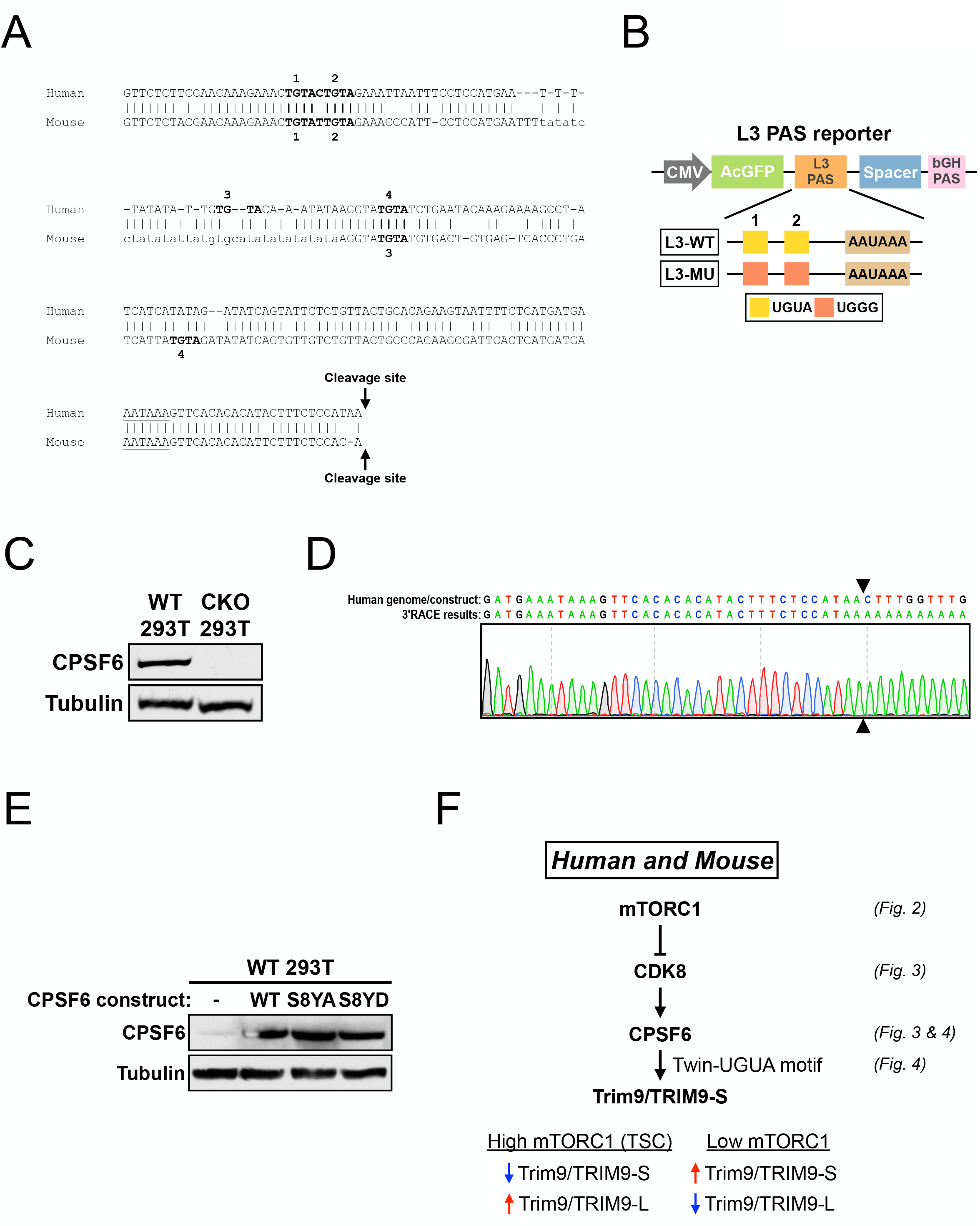
Characterization of reagents and tools for the PAS reporter assay, related to Figure 4. (**A**) Pairwise genomic sequence alignment (0~-200bp; 0 = cleavage site) between human TRIM9-S PAS and mouse Trim9-S PAS. TGTA motifs are numbered from 5’ to 3’ and are shown in bold. The poly(A) signal, AATAAA, is underlined. See Fig. 4A for the illustrated version. (**B**) Illustrations showing the design of wildtype (L3-WT) and mutant (L3-MU) L3 PAS reporters. (**C**) Western blots showing *CPSF6* protein expression in wildtype (WT) and CPSF6-KO (CKO) 293T cells. Tubulin: loading control. (**D**) Sanger sequencing results from a 3’ RACE experiment in 293T cells showing the actual cleavage site (indicated by arrowheads) used in the TRIM9-S WT PAS reporter. Also see (A) for the endogenous cleavage site of TRIM9-S. (**E**) Western blots showing *CPSF6* protein expression in untransfected 293T cells or 293T cells transfected with different CPSF6 constructs (WT, S8YA, and S8YD). Tubulin: loading control. (**F**) Illustrations summarizing how mTORC1 regulates TRIM9-S expression in human and mouse in addition to how high/low mTORC1 activities change *Trim9/TRIM9* isoform expression. Relevant figures are listed on the right.

**Figure S5.**
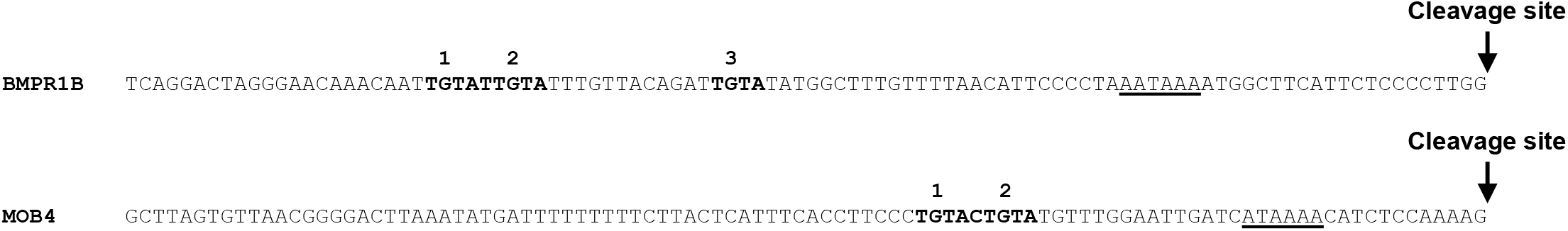
Genomic sequence (0~-100bp; 0 = cleavage site) of human *BMPR1B* and *MOB4* distal PASs, related to Figure 5. TGTA motifs are numbered from 5’ to 3’ and are shown in bold. The poly(A) signal, AATAAA, and a putative non-canonical poly(A) signal, ATAAAA, are underlined. See Fig. 5E for the illustrated version.

**Figure S6.**
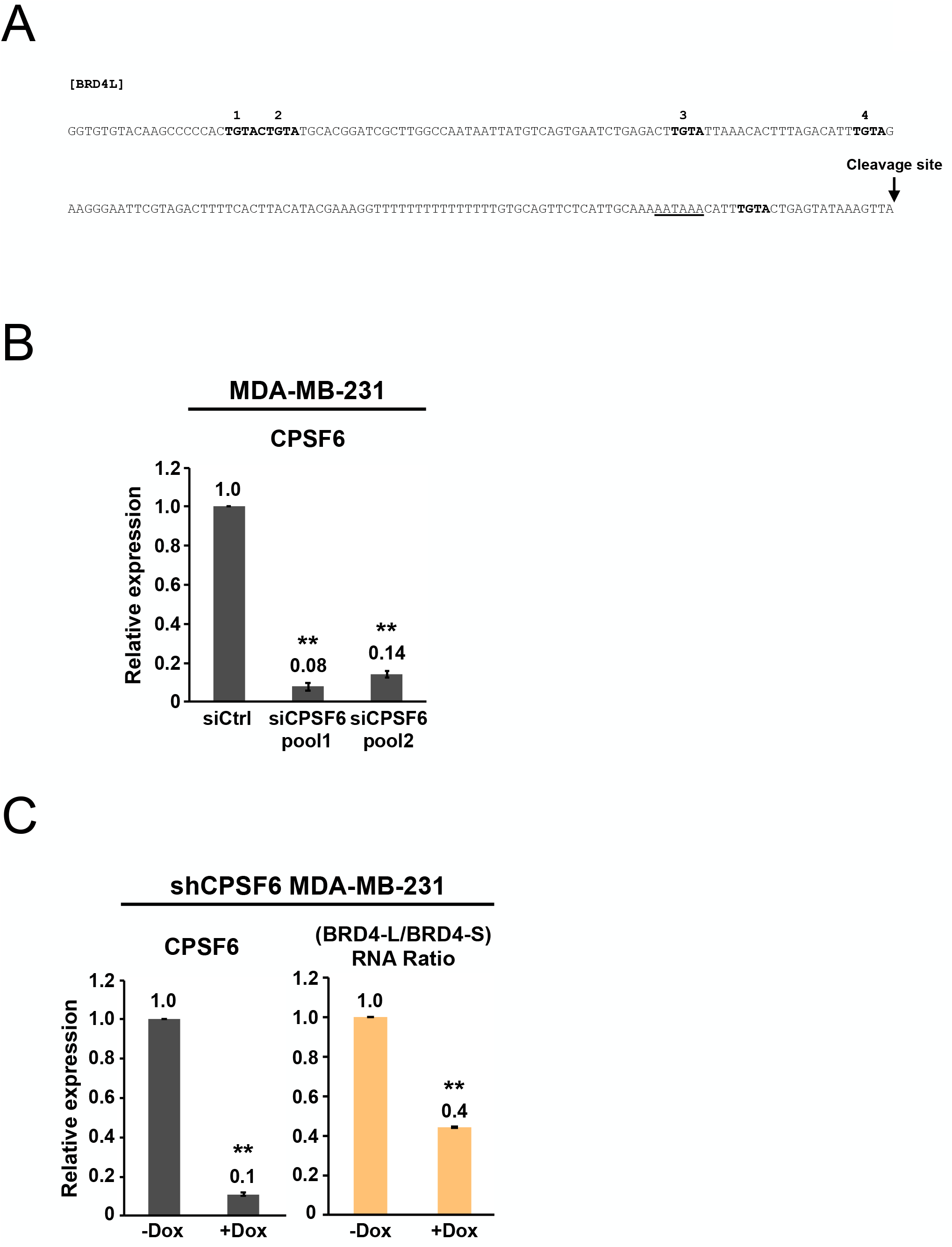
*CPSF6* knockdown in MDA-MB-231 cells, related to Figure 6. (**A**) Genomic sequence (0~-200bp; 0 = cleavage site) of human BRD4-L PAS. TGTA motifs are shown in bold. The 3 TGTA motifs preceding the poly(A) signal are numbered from 5’ to 3’. The poly(A) signal, AATAAA, is underlined. See Fig. 6A for the illustrated version. (**B**) Bar graphs showing the knockdown efficiency of *CPSF6* siRNAs measured by RT-qPCR 72 h after transfection in MDA-MB-231 cells in the same experiments shown in Fig. 6F (n=3). (**C**) Bar graphs showing expression of *CPSF6* mRNA (left) and the ratio of *BRD4* mRNA isoforms (right) measured by RT-qPCR in shCPSF6 MDA-MB-231 cells with (+Dox) and without (-Dox) doxycycline treatment for 72 h from 3 independent experiments (n=3). Dox: 1 μg/mL doxycycline. Error bars indicate SEM. Statistical significance is determined by two-tailed t-test. **: p<0.01.

**Figure S7.**
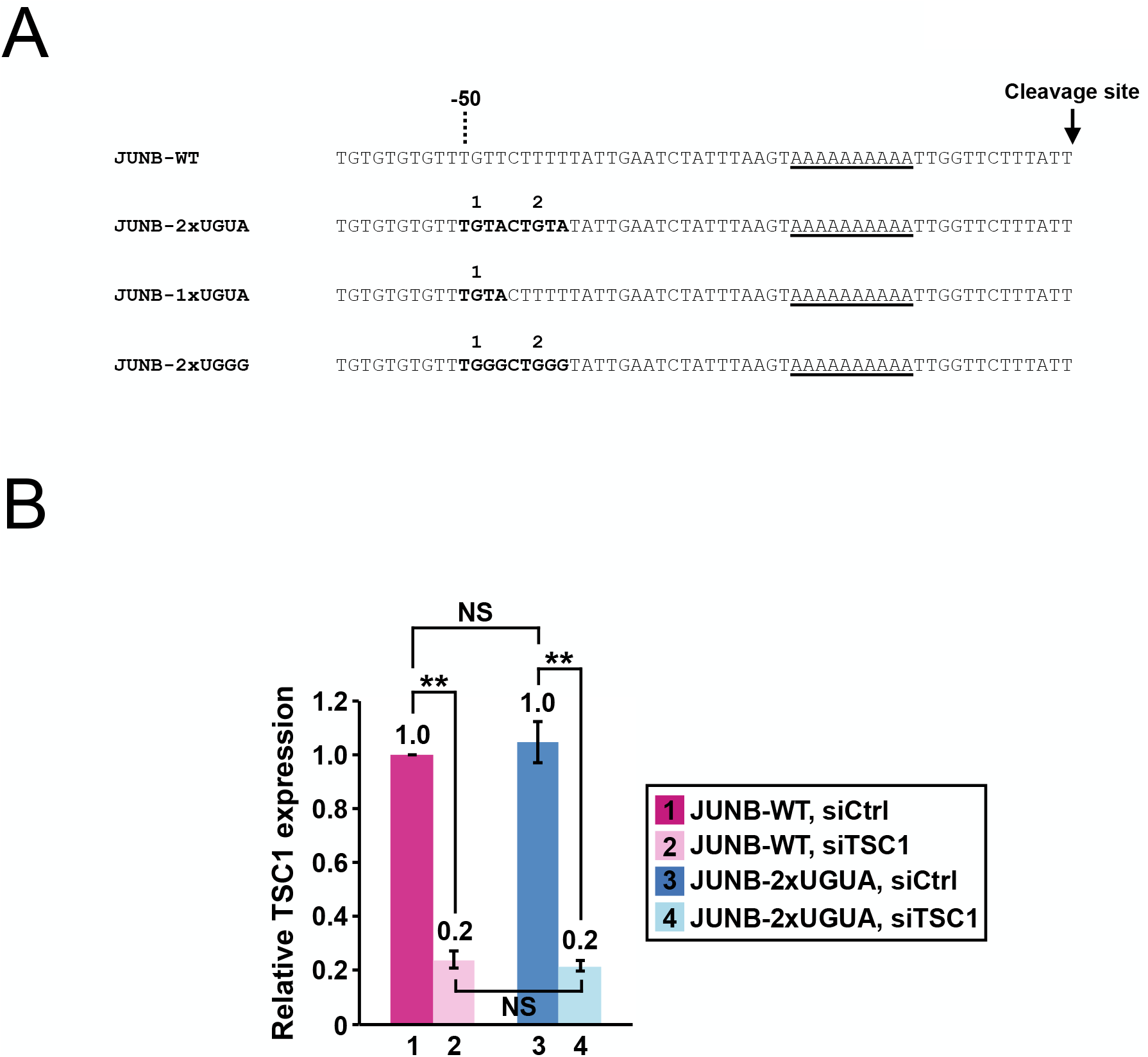
Nucleotide sequences of wildtype and modified *JUNB* PASs for the reporter assay, related to Figure 7. (**A**) Nucleotide sequences of wildtype and modified human *JUNB* PAS. TGTA and TGGG motifs are numbered from 5’ to 3’ and are shown in bold. An adenine-rich element containing the putative non-canonical poly(A) signal is underlined. See Fig. 7A for the illustrated version. (**B**) Bar graphs showing *TSC1* knockdown efficiency measured by RT-qPCR in HeLa cells in the same experiments shown in Fig. 7D (n=3). All expression levels are relative to group 1 (JUNB-WT reporter, siCtrl transfection). Error bars indicate SEM. Statistical significance is determined by two-tailed t-test. NS: not significant, **: p<0.01.

